# Evolution of the Batoidea Pectoral Fin Skeleton: Convergence, Modularity, and Integration Driving Disparity Trends

**DOI:** 10.1101/2024.06.26.600866

**Authors:** Faviel A. López-Romero, Eduardo Villalobos-Segura, Julia Türtscher, Fidji Berio, Sebastian Stumpf, Richard P. Dearden, Jürgen Kriwet, Ernesto Maldonado

**Affiliations:** EvoDevo Research Group, Unidad de Sistemas Arrecifales, Instituto de Ciencias del Mar y Limnología, Universidad Nacional Autónoma de México, Puerto Morelos, Quintana Roo, México; University of Vienna, Faculty of Earth Sciences, Geography and Astronomy, Department of Palaeontology, Evolutionary Morphology Research Group, Josef-Holaubek-Platz 2, 1190, Vienna, Austria; University of Vienna, Vienna Doctoral School of Ecology and Evolution (VDSEE), Djerassiplatz 1, 1030, Vienna, Austria; Department of Zoology, Stockholm University, Svante Arrhenius väg 18B, 114 18, Stockholm, Sweden; Vertebrate Evolution, Development, and Ecology, Naturalis Biodiversity Center, Darwinweg 2, Leiden, 2333 CR, The Netherlands; University of Birmingham, School of Geography, Earth & Environmental Sciences, University of Birmingham, Edgbaston, Birmingham B15 2TT, UK

**Keywords:** Batoidea, Pectoral fin skeleton, Evolution, Modularity, Disparity, Convergence

## Abstract

Batoids (skates and rays) are the most speciose group of cartilaginous fishes. Their body plan represents diverse ecologies and swimming modes. Early skeletal fossil remains, and recent phylogenetic analyses suggest that convergence has occurred within the batoids several times independently. The drivers for such disparity patterns and possible association with modularity and phenotypic integration among batoids are not fully understood. Here we used geometric morphometrics and phylogenetic comparative methods to characterize the evolutionary trends of the basal fin skeleton of batoids and sharks. Our analyses show that the morphological variation has a strong phylogenetic signal. Interestingly, the most speciose orders of batoids display low morphological disparity. Reef and freshwater species, show increased evolutionary rates. Meanwhile, the swimming mode shows different rates depending on the fin structure analyzed. A higher modularity and integration signal suggest that the pectoral fin of batoids has experienced mosaic evolution. The low morphological disparity might be associated with high integration. We find support for convergence between Jurassic, Cretaceous, and Extant guitarfishes, however, not completely between sharks and batoids. Our findings suggest that habitats and swimming mode have shaped the pectoral fin evolution among batoids, and at the same time batoids have constrained their basal fin skeleton.

## 1. Introduction

Rays, skates and guitarfishes (hereafter “batoids”) comprise the most speciose group of cartilaginous fishes, with nearly 620 species described to date (1, 2). Batoids have diversified in virtually all possible aquatic environments, from open ocean to freshwater and from nearshore reefs to the deep sea (3, 4). The most striking feature of batoids is their dorsoventrally flattened body, expanded pectoral fins to form a closed disc, which display a high diversity of shapes (5, 6, 7) (Figure 1A). The basic structure of the pectoral fin is composed of three basal elements which articulate to the coracoid bar by condyles (Figure 1A and B). The wing-like fins also show several modifications, the radials showcase differences in the mineralization associated with the swimming type (8). Batoids have a remarkable long fossil record with several groups represented by completely articulated specimens (9). The earliest remains of batoids are found in the Early Jurassic and several articulated specimens in the Middle Jurassic (10, 11, 12, 13). Extinct representatives of modern groups of batoids showcase a wide morphological disparity in several traits which are interpreted as a mixture of plesiomorphic and derived features (14, 15, 16, 17). Meanwhile, modern batoids diversified around the Lower Cretaceous, making them a long-standing group that has endured extinction events (11, 13, 18, 19). This provides a unique opportunity to study the evolutionary trends associated to the pectoral fin in deep time.

**Figure 1.**
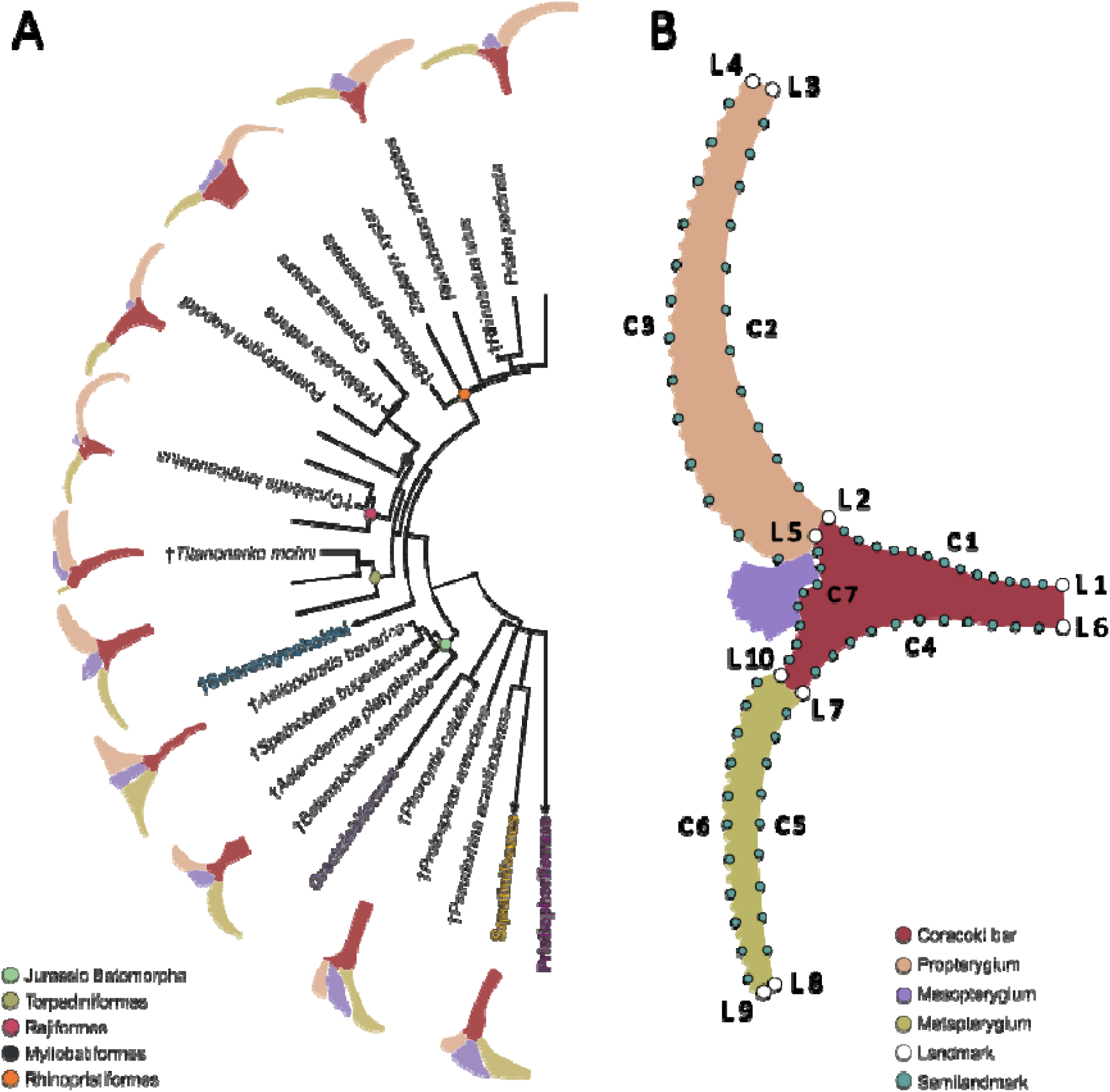
Simplified phylogeny of the studied group of elasmobranchs displaying the corresponding skeletal anatomy of the pectoral fin (A). Landmark coordinates scheme followed to perform the statistical shape analysis (L1-10 = fixed landmarks; C1-7 = Curve landmarks) (B).

A dorso-ventrally flattened body has occurred independently in cartilaginous fishes, like in several Paleozoic forms, holocephalans, and Mesozoic to modern sharks (20, 21, 22 5, 23). As a total group, batoids are estimated to have evolved during the Permian at the split with their sister group, the sharks (24). Batoids have acquired highly disparate body forms, like the sawfish and even the bowmouth guitarfish with a shark-like shape (25). Consequently, the resolution of the relationships within the main four orders (Rajiformes, Myliobatiformes, Torpediniformes and Rhinopristiformes) becomes relevant to assessing the patterns of morphological disparity. Even the relationships with their sister group (sharks) presented different arrangements through the years (26, 27, 28, 29, 30, 31, 32, 33, 34, 35). The outline shape of the pectoral fin seems to reflect some relation to their phylogenetic relationships. Previous studies indicate that highly specialized groups like stingrays display a high morphological disparity (6, 7), which is explained by their swimming mode in terms of the aspect ratio of the fins (the area supported by plesodic radials) (6). However, the internal features can reveal differences in disparity among sharks, which are not necessarily under the same pressure (36). This highlights the importance of assessing the anatomical features, to understand the underlying processes leading to convergence patterns. Especially considering how the skeletal elements of the pectoral fin are articulated to each other, might indicate a possible constraint imposed by their body plan.

The extent to which distinct components of anatomical features are linked to each other, and the patterns of covariation they display is known as phenotypic integration (37). These patterns can indicate if two or more structures covary between them to consider them as a composed unit (module) (37). The extent of integration within a group has been associated to an increase or decrease in the amount of morphological disparity (38). Patterns of variation representing a constraint due to a higher pattern of integration can lead to the evolution of extreme shapes along a single axis of variation (39, 40). Besides, highly integrated phenotypes often tend to display low morphological disparity. In this sense, the variation is constrained although this does not prevent the group from displaying differences in species diversity (41). Therefore, there can be groups with a high number of species but with low disparity, and the opposite case can also occur (42). Together, this can indicate trends through the history of taxonomic groups in deep time, like the release of disparity following an event in the past and the related macroevolutionary consequences (43). The patterns of modularity and integration are relevant because the traits can evolve in different trajectories, leading to specialized forms (44). Among elasmobranchs, it has been shown that batoids display higher modularity signal in the skull than sharks, which might have facilitated their diversification driven by the prey-acquisition strategies (45). However, little is known about the possible role of modularity and integration in the evolution of the pectoral fin, especially in relation to swimming.

Our study has four main objectives. First, to explore the morphological variation of the skeletal elements composing the base of the pectoral fin among extant and extinct batoids to determine patterns of disparity. Second, to investigate if individual elements of the pectoral fin experienced differences in morphological evolutionary rates in relation to swimming type and habitat. Third, we seek to find whether modularity or integration has contributed to the shape variation of the pectoral fins among sharks and batoids. Finally, to determine the patterns of convergence between sharks and batoids, and within batoids. Altogether, our study aims to understand the drivers of the pectoral fin shape evolution in the major group of cartilaginous fishes.

## 2. Material and Methods

### (a) Data acquisition

We gathered information of the internal anatomy of batoids with several radiographs taken from available depositories in iDigiBio. The depositories contain information from the following museums: Smithsonian Institution, National Museum of Natural History (USNM), Natural History Museum, London (NHMUK), Museum national d’Histoire naturelle (MNHN), Australian Museum (AM), California Academy of Sciences (CAS), Florida Museum of Natural History (FLMNH), Harvard University, Museum of Comparative Zoology (MCZ), Naturhistorisches Museum Wien (NMW), Field Museum of Natural History (FMNH). Besides, segmentations from CT-Scans were performed with the software Slicer3D (v. 5.2.2), Mimics (v. 23.0) (Materialise), and screenshots of the 3D images were taken in ventral view. The CT-Scans are available from FigShare (46), Morphosource, and Chondrichthyan Tree of Life. Additionally, photos of fossil specimens come from museum collections: Swedish Museum of Natural History (NRM), Staatliches Museum für Naturkunde Stuttgart (STU), NHMUK, MNHN, AMNH). A total of 362 specimens from which 330 specimens belonging to batoids, representing 194 species. We verified the taxonomic assignment in the Eschmeyer Catalogue (2) and FishBase (47). From FishBase, we obtained information regarding depth distribution and environmental occurrences for each species. For fossil species, we obtained information regarding the environment with the Paleobiology Database (48) and from literature.

### (b) Phylogenetic reconstruction

We examined the phylogenetic relationships of the various elasmobranch taxa using a modified data matrix of Villalobos-Segura et al. (35), to provide a phylogenetic context for the macroevolutionary analyses. The only modifications made to the matrix were in the number of terminals, to accommodate the increased number of taxa included in the current analysis. The matrix includes fossil chondrichthyans from the Paleozoic, Mesozoic, Cenozoic, and recent taxa. The fossil genus †*Doliodus* served to root the phylogenetic analysis. Two positive constraints were enforced, one for the whole Batoids to ensure that no wild card taxa fell outside this group. This constrain is not necessary as the analysis can be carried out without it, since there were no batoid taxa falling out of this group. Another constraint was inflected on the Torpediniformes, to include all molecular groupings at the order level, to accommodate the inclusion of the recent taxa, without radically increasing the number of characters and performing an extensive anatomical study, which would be beyond the scope of the present study, (see electronic supplementary material) (30, 25, 4). The remaining phylogenetic associations were left unconstrained, to ensure reflecting the phylogenetic uncertainty associated to the morphological characters and the discrepancy between the phylogenetic hypotheses under morphological and molecular data. The resulting data matrix was assembled in Mesquite (v. 3.81) (49) and contains 253 terminals and 143 characters (see electronic supplementary material).

A parsimony analysis was conducted using TNT (v. 1.6) (50). A new technology search was performed with 1,000 ratchet iterations, TBR (tree bisection and reconnection) and SPR (subtree pruning and regrafting) were used as the algorithm for branch permutations, holding one tree, additionally ten cycles of Tree drifting. This search was performed until 10 hits of the minimum score tree were reached. This search protocol was run ten times, saving the trees found on each search. All ten searches recovered the same strict consensus (see electronic supplementary material), suggesting an adequate search of the tree space. All the most parsimonious trees recovered in these ten searches were kept, but only the trees with unique topology were later used in the macroevolutionary studies. Tree branch lengths and likelihood scores were calculated using PAUP (v. 4.0a) (51) under the Mkv model with the gamma rate parameter, following the approach used by Brazeau et al. (52).

### (c) Geometric Morphometrics

A landmark configuration was used to describe the shape of the pectoral fin (Figure 1B, supplementary table 1). We considered the coracoid bar, the first element of the propterygium, and the first element of the metapterygium. The mesopterygium was excluded from the analysis since it is not always present, or it is fused with the radials. Only the first element of the propterygium and metapterygium, respectively, were used for comparison, because the number of elements is variable between species. Only the first element is consistently present allowing the assumption that these elements are homologous across taxa. The 2D landmarks and semilandmarks coordinates were captured with TPSDig2 (v. 2.31) (53). The coordinates were then subject to a Generalized Procrustes Analysis (GPA) using the bending energy to slide the semilandmarks (54, 55). This was performed with the gpagen function from the R package geomorph (v. 4.0.7) (56). Coordinates partitions of each element were also subjected to a GPA, because we intended to trace shape changes of each individual element. The aligned coordinates were then used to perform a principal component analysis to visualize the variation among the individuals and explore the shape changes. This was performed for the full configuration and each element separately with the gm.prcomp function in geomorph. The original coordinates were then averaged to the species level and used with the phylogenetic hypothesis including fossil species to perform a phylogenetic aligned component analysis (PACA) (57) which aligns phenotypic data to the phylogenetic signal.

### (d) Morphological Disparity

We estimated the disparity per group using the sum of variances with all landmark’s configurations. We used the package DispRity (v. 1.8) (58) and divided the set into the different taxonomic groups displayed in Figure 1 (Rajiformes, Rhinopristiformes, Myliobatiformes, Torpediniformes, Squatiniformes, Orectolobiformes, Pristiophoriformes, Jurassic Batoids, Cretaceous Rhinopristiformes, sclerorhynchids, Cretaceous Rajiformes, and Eocene Myliobatiformes). We also considered the disparity of other grouping factors. We used a modified classification by Martinez et al (59) to compare between “deep sea”, “shelf”, “reef”, and “freshwater” occurring species. We compared the assigned swimming types into the categories “undulatory”, “oscillatory” (mostly present in Gymnuridae and Myliobatidae, and Mobulidae), and “axial undulatory”, since the pectoral fin is also linked to a swimming type (60, 8). We used the calibrated phylogeny and obtained the ancestral states for the nodes in the PACA. Together with this matrix and the phylogenetically aligned components (PAC) that explain 99% of the variation, we performed a disparity through time analysis using the sum of variances to observe changes throughout the history of batoids and sharks that could be associated with past geological events.

### (e) Phylogenetic comparative methods

We used a subset of 100 random trees from the 420 trees obtained to perform the phylogenetic comparative methods to account for phylogenetic uncertainty. We trimmed the phylogeny to contain only the taxa used. We calibrated the tree with information about the first appearance in the fossil record for each group and the fossils used. We used the scaleTree function in RRphylo package (v. 2.8) (61). We used the categories of habitat occupancy and swimming type to estimate the discrete evolutionary rates. First, we compared the support of the fitting of the Equal Rates, Symmetric, and All Rates Different models using the AIC and logLikelihod to evaluate the support using the fitMk function in castor (v. 1.8) (62). With the selected model, we used the sim.map function in phytools (v. 2.1-1) (63) to trace the history of traits. We performed the mapping on the 100 random trees and used these trees for the following analysis. From the PACA, we selected the components explaining up to 99% of the variation, these components were then used with the mvgls function in mvMorph (v. 1.1.9) (64) to obtain the morphological evolutionary rates of the shape variables conditioned to the discrete traits. Additionally, we used these selected components with the mvgls, followed by the manova.gls function to estimate the association of shape with the categorical variables using Pillai’s test for significance. We performed this analysis on a set with only extant batoids.

We also used another approach to estimate the rates independently of a model. We selected the phylogeny with the best score and calibrated the tree with information from the literature. With the calibrated tree we used the RRphylo function from the RRphylo package (v. 2.8) (61) to perform a phylogenetic ridge regression and obtain the rates per branch. With the search.shift function we compared the rates of the nodes of each group of interest (the main orders of batoids) to compare if each different taxonomic group has experienced a shift in the rates. We also estimated the shift rates for each clade with the RRphylo search.shift function and verified the consistency of the results using 100 random trees with the overfitRR function. Because several instances of convergence have been suggested between groups of sharks and batoids and within groups of batoids, we performed a convergence test in RRphylo with the search.conv (65). We took the Procrustes distances of the aligned coordinates and performed a cluster analysis using the UPGMA method. We then used this distance dendrogram with the phylogeny with the function cophylo from phytools to visualize the groups which converge into a cluster (Supplementary Figure 1). We compared the resulting tangled groups to test for convergence. We performed other comparisons specific to groups like sclerorhynchids, Squatiniformes, Pristiophoriformes, and Pristidae. We performed a specific node search for convergence with these groups as well as between selachians and batoids to further explore a possible convergence often mentioned between angel sharks and batoids.

### (f) Modularity and Integration

Finally, the coordinates configurations were subdivided into five possible module hypotheses (Supplementary Figure 2) considering: each element independent from each other (H1), a configuration with the propterygium and pectoral girdle as a module (H2), a configuration of the coracoid bar and metapterygium as a module (H2), a configuration of propterygium and metapterygium as a module separated from the coracoid bar (H3), and a null hypothesis of no modularity (H0). We compared the possible hypotheses with the compare.phylo.CR function in geomorph (66), which considers the covariance and the effect size to determine the strength of a modularity signal. The selected hypothesis, based on the covariance ratio effect sizes (Zcr), was used to compare the modularity signal by groups. We divided the subsets into different taxonomic orders of extant batoids, another comparison between extant and extinct batoids and a comparison of batoids and sharks. We estimated integration with two alternative methods. First, we used the partial least squares approach (67) with the defined modularity hypothesis from the previous analysis. We used another approach (68) to assess the pattern of local or global integration changes in terms of self-similarity (no interpretable structure at any temporal scale). We compared the same groups as the ones for the modularity test.

## 3. Results

### (a) Pectoral fin shape variation

The results from the Principal Component Analysis are similar to the ones derived from the phylogenetic aligned component analysis (PACA) (Supplementary Figures 3-6). The PACA shows that in the first component (PAC1) (92.42% of the variation) shape changes in the negative scores are associated with elongation of both propterygium and metapterygium, a short coracoid bar along the lateral axis and elongated in the joints with the propterygium and metapterygium (Figure 2A). Most of the observed species in these scores correspond to Myliobatiformes and the extinct Rajiformes genus †*Cyclobatis*. In the positive scores, the coracoid bar extends laterally from the midline to the propterygium and metapterygium articulations becoming slender. The sharks (Orectolobidae, Squatiniformes, and Pristiophoriformes) are located on these positive scores, without any overlap from a batoid species, only a sclerorhynchid (†*Ptychotrygon*) and a fossil torpediniform (†*Titanonarke*) seem to display a similar shape. Most of the species placed towards the mean values correspond to Rhinopristiformes, both extinct and extant. Jurassic batoids are overlapped by Rhinopristiformes in the morphospace. Regarding PAC2 (6.04% of the variation), the minimum values show a medially short and posteriorly elongated coracoid bar where the metapterygium articulates. The propterygium is notably reduced on these scores while the metapterygium is quite elongated. All the species found here correspond to modern Rajiformes. On the positive PAC2 scores, the shape changes correspond to a medially elongated coracoid bar and a relatively short distance on the articular condyles, resembling the sharks’ coracoid bar. The propterygium is elongated, however, it is not quite as long as in Myliobatiformes, whereas the metapterygium is shorter than the rest of the other ones in the morphospace. Overall, the positive PAC2 scores correspond to Torpediniformes (electric rays), which tend to display unfused coracoid bar on both antimeres and a notable reduction of the metapterygium. All the main orders of batoids are separated rather consistently in the morphospace when we consider the three structures at the same time in the analysis.

**Figure 2.**
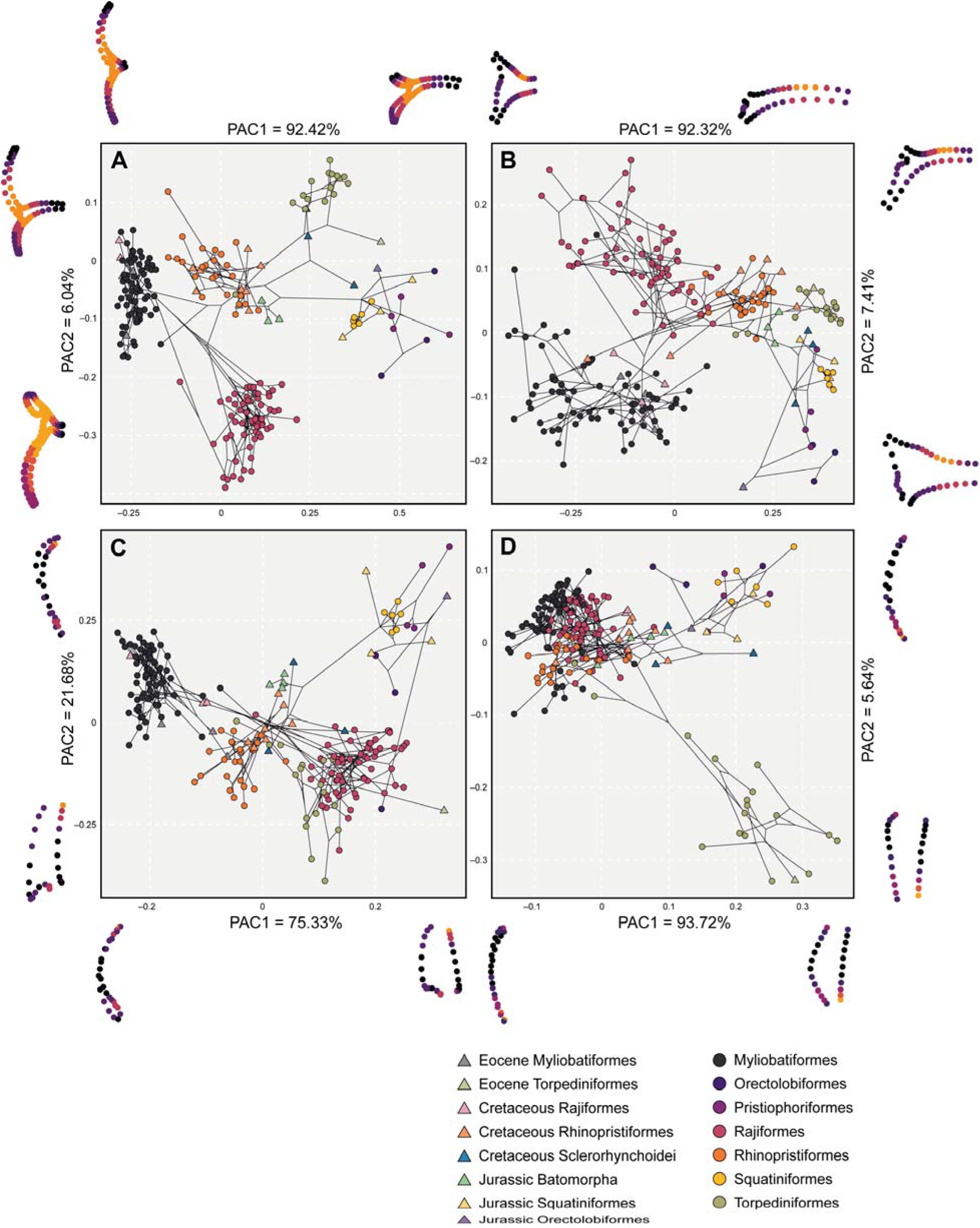
Phylomorphospaces for each landmark configuration, displaying averaged specimens at species level. Symbols on the right indicate the groups of each taxonomic order as circles and extinct groups as triangles. On each side of each axis, we show the extreme shapes for each component. A) Full landmark configuration, B) Coracoid bar configuration; C) Propterygium configuration, and D) Metapterygium configuration.

When each structure is analyzed separately, we observe a different pattern. The coracoid bar still represents a rather good trait for distinction and is consistent with the main groups (Figure 2B). The PAC1 (92.32% of the variation) is explained by lateral elongation of the bar on the positive scores and narrow distance on the articulation region. The negative scores the bar is short and wide (antero-posteriorly) in the articulation area. Myliobatiformes and Rajiformes are distributed in a gradient-like pattern along these scores, followed by Rhinopristoformes and at the extreme the Torpediniformes and finally sharks in the positive scores. The second PAC2 (7.41% of the variation) explains mostly the elongated posterior articulation region of the coracoid bar, where the metapterygium attaches and a slender bar, Rajiformes are located in these scores. The negative scores show a wide coracoid bar, especially in the anterior part as seen in most members of Myliobatiformes like Gymnuridae and Rhinobatidae. The propterygium PACA (Figure 2C) displays a relatively good separation among groups. The propterygium is short on the negative scores of the PAC1 (75.33% of the variation) with a wide articular facet, strongly curved inwards with an anterior elongation which tapers. The shapes found on the positive scores correspond to sharks which have an overall short propterygium, these are followed towards the negative score by the Rajiformes and Torpediniformes, which overlap in this part of the morphospace. Meanwhile, the Jurassic batoids, sclerorhynchids, and Rhinopristiformes overlap in the middle. Myliobatiformes is located on the negative extreme values. The PAC2 (21.68% of the variation) on the positive scores shows a rather bulky anterior portion of the propterygium that tappers in the articular region. In the negative scores, it becomes more rectangular in shape with a narrow articular region. The extreme positive PAC2 shape is not usually seen among any group, although it can be similar in some fossil shark forms like extinct †*Pseudorhina*, although this one has a wider articulation region. The shapes in the negative scores correspond to members of the Torpediniformes. Finally, the metapterygium (Figure 2D) shows a rather unique pattern with an overlap of Rajiformes, Myliobatiformes, and Rhinopristiformes, along with fossil forms. The PAC1 (93.72% of the variation) explains changes in the posterior elongation of the metapterygium. The positive scores display a rather short and curved metapterygium, while the metapterygium is slender and posteriorly elongated in the negative scores. Only Torpediniformes displays a short metapterygium and this is indicated as a divergence in shape from the main cluster. The PAC2 (5.64%) shows a similar pattern with the positive scores expressing an elongated and curved metapterygium, while the negative scores typify a short and straight on the outer side metapterygium.

### (b) Phylogenetic signal

Our results suggest that the strongest separation between groups occurs when all structures are considered. However, both the coracoid bar and propterygium show a consistent separation between groups with some overlap. This suggests that the phylogenetic signal also varies for each structure conversely to analyzing all three structures together. Indeed, the phylogenetic signal for the whole configuration indicates that there is a signal mostly separating clades (λ = 0.9696; Kmult = 1.5387), while this signal is reduced with more internal variation within the clades when the structures are analyzed separately. In this regard, the coracoid bar still displays a high phylogenetic signal (λ = 0.8564; Kmult = 0.7864), followed by the metapterygium (λ = 0.9223; Kmult = 0.7585) and propterygium (λ = 0.8752; Kmult = 0.6262).

### (c) Morphological disparity by groups and through time

We estimated the disparity of the different groups to discern whether there are differences between extant and extinct counterparts, as well as different habitat groups and swimming types groups. When considering the full configuration of landmarks, the estimated Procrustes variance for the morphological disparity shows that Torpediniformes displays the highest disparity among all the groups. They are followed by Orectolobiformes (*Orectolobus* spp, *Eucrossorhinus*, and *Brachaelurus*), although they are represented by only few specimens and do not show the full disparity extent among Orectolobiformes. Squatiniformes displays the lowest disparity among the extant groups (Figure 3A). Interestingly, despite being the most speciose, both Myliobatiformes and Rajiformes display low disparities (Figure 3A). The extinct sclerorhynchids have the highest disparity among them all, followed by Cretaceous Rhinopristiformes. The Jurassic batoids, conversely, display low disparity. Overall, the extinct taxa do not display higher disparity than extant groups, except for the sclerorhynchids in either the full configuration or when isolated elements are considered (Supplementary Table 2). In view of this pattern, we explored a temporal component to reveal the disparity through time. For both the complete set and the set with batoids, there only is a pattern of gradual increase in disparity (Figure 3B; Supplementary Figure 7) starting in the Jurassic when the first holomorphic batoids appear in the fossil record. There is a steady increase in disparity until the Cretaceous when it reaches a maximum, followed by a sudden decline during the KPg extinction event. This reduction of disparity is followed by a constant increase from the Eocene persisting until today, suggesting a recovery delay of disparity right after the KPg extinction event. The disparity through time for each landmark configuration shows a very similar pattern as the one shown for the whole landmark configuration (Supplementary Figure 8). The disparity of habitat occupancy shows the highest disparity occurs among the reef and shelf associated species (Figure 4A and Supp. Table 2). Deep-sea species display an overall low disparity, most species in this category are Rajiforms, which already display low disparities. Finally, the freshwater groups display the lowest disparity among all the groups. This category includes only members of the Potamotrygonidae (Myliobatiformes), which are highly specialized batoids in respect to their life-style. The comparison of each individual element and the whole configuration displays very similar results for every habitat category (Figure 4A, Supplementary Table 2). Regarding the swimming type, we found that species using axial-undulatory display the highest disparity, followed by the undulatory, and finally oscillatory type (Figure 4B). In the case of the oscillatory swimming type, this might be expected because only families of Myliobatiformes are in this category (Aetobatidae, Gymnuridae, Myliobatidae, and Mobulidae) (Supplementary Table 2). However, analyzing the elements separately, we observe that the disparity of the coracoid bar in the undulatory swimming species is higher than the other two categories. Disparity of the propterygium is higher among undulatory swimming species, followed by axial undulatory species. Finally, the disparity of the metapterygium indicates that the axial undulatory type is higher than the other two categories (Supplementary Table 2). Considering the pairwise comparisons for all examined groups, we found that for most of the cases a clear difference in their disparity regardless of the subdivision of the compared landmarks or as a whole configuration including all the elements (Supplementary Tables 3-6).

**Figure 3.**
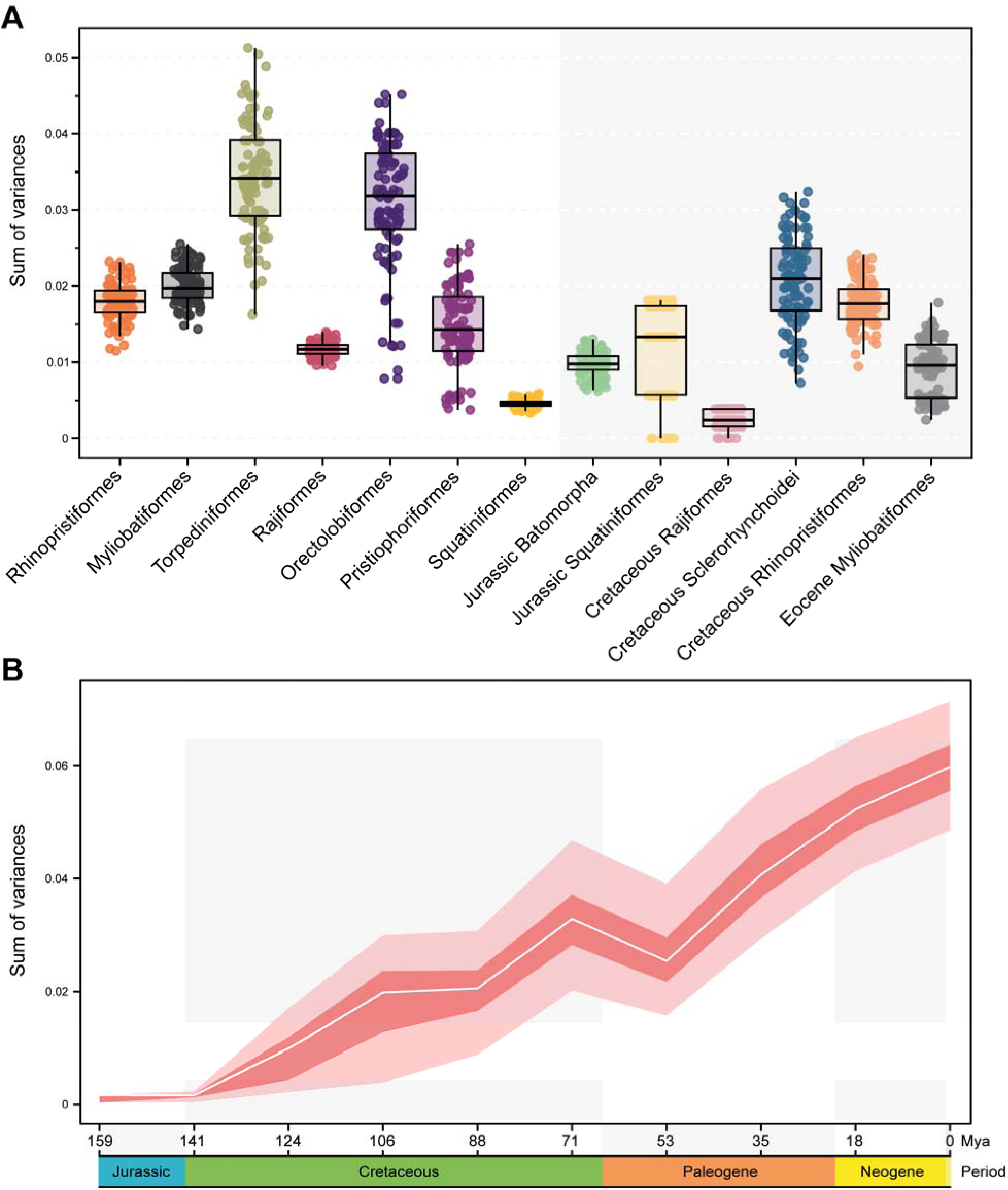
Morphological disparity estimated as the sum of variance by taxonomic groups (A), through time (B).

**Figure 4.**
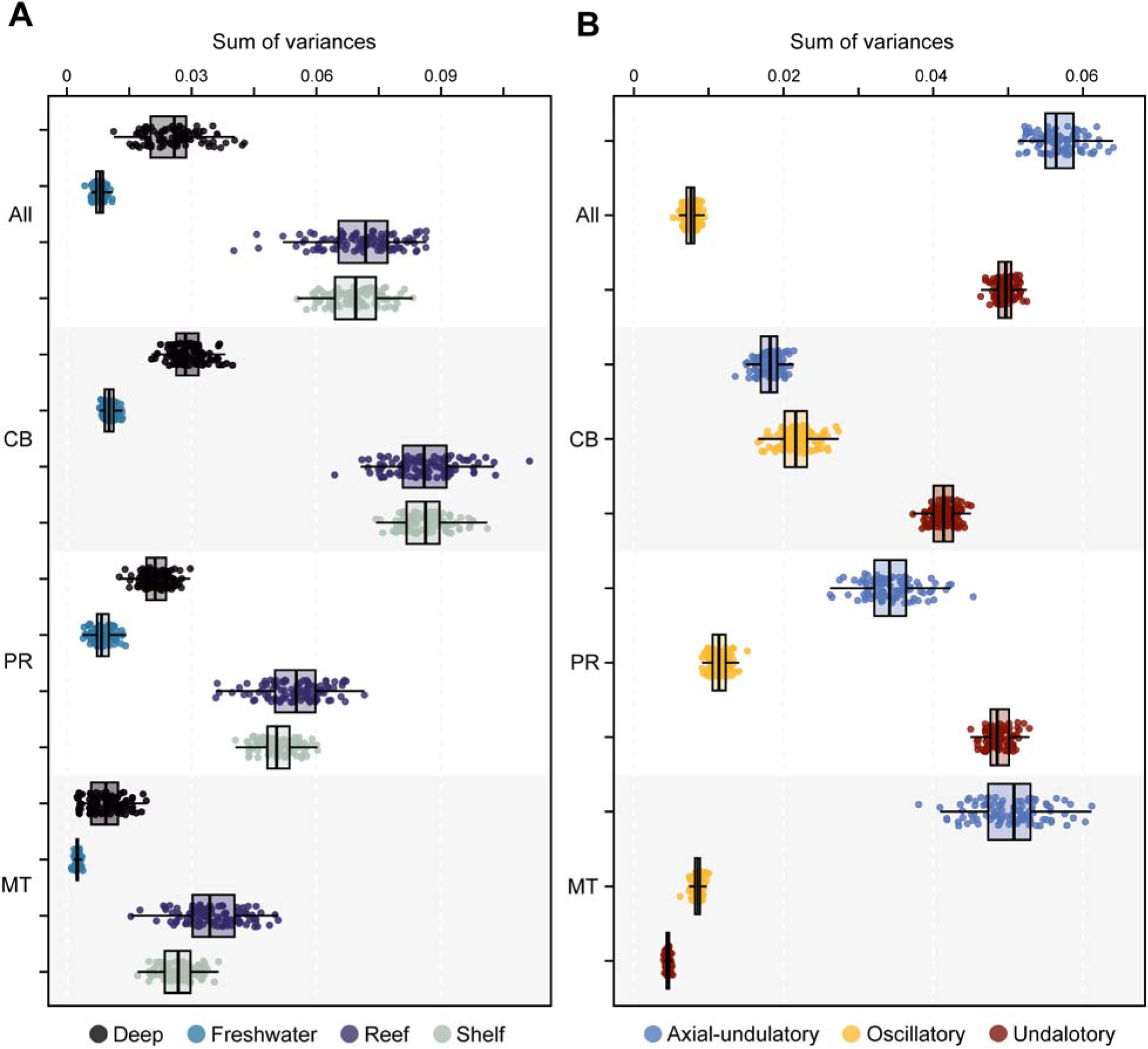
Morphological disparity estimated as the sum of variance by habitat groups (A), and by swimming type (B). All: All elements; CB: Coracoid Bar; PR: Propterygium; MT: Metapterygium.

### (d) Evolutionary rates

Subsequently, we investigated whether the morphological evolutionary rates display a shift in the phylogeny when considering the shape variables. The results indicate a shift in rates in only a few batoid orders (Figure 5, Supplementary Table 7). The sclerorhynchids are among the extinct taxa that display a positive shift in rates, and among extant taxa, there seems to be a positive shift in Rajiformes, whereas Myliobatiformes displays a significant negative shift when the full configuration is considered. In the case of the coracoid bar, we find that almost all batoid clades experienced a positive shift, while selachians do not appear to display a significant shift (Supplementary Figures 9-11). The rates for both Myliobatiformes and Rhinopristiformes appear to be higher than for the other orders. For the propterygium, we found that only Jurassic batoids and selachians do not display a significant shift in the rates. However, there is a significant shift at the basal node of all batoids. Similarly, the metapterygium displays a pattern of positive shifts in the examined main clades, while selachians do not display a significant shift at all. All these results present a consistent pattern when accounting for phylogenetic uncertainty (Supplementary Tables 7 and 8), with 100% of all the instances finding the same shift for the analyzed clades.

**Figure 5.**
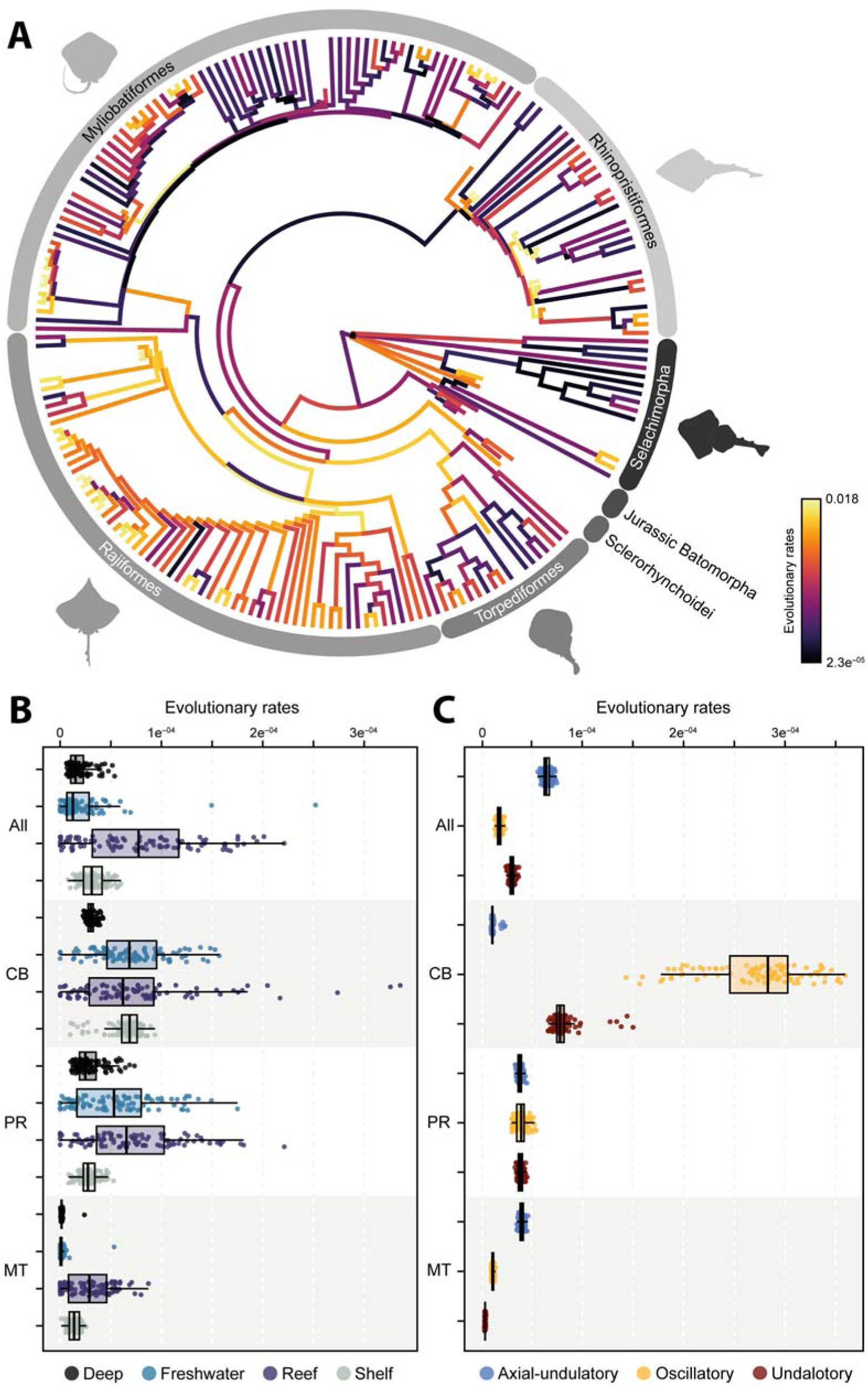
Morphological evolutionary rates between each taxonomic group (A), Habitat (B), and Swimming type (C). The values on the tree are expressed as absolute rates from the tips after ridge arch regression. The rates for the two remaining groups are expressed as sigma σ^2^. All: All elements; CB: Coracoid Bar; PR: Propterygium; MT: Metapterygium.

The evolutionary rates expressed as σ^2^ for the swimming type and habitat occupancy performed on the extant set, indicates that for the whole landmark configuration, shelf species tend to evolve faster, although the effect is not significant (Supplementary Tables 9-10). The coracoid bar shows a significant effect, with the reef species displaying high evolutionary rates, followed by freshwater species (Figure 5B). This is similar to the results for the propterygium, which shows a significant effect (Supplementary Tables 9-10). In the case of the metapterygium, the habitat does not appear to have a significant effect on the shape, although reef and shelf species display higher rates than deep sea and freshwater species. Regarding the swimming type, we find that, considering the whole configuration, the axial undulatory type evolved faster than the undulatory and oscillatory types (Figure 5C and Supplementary Tables 9-10). However, for the coracoid bar, the oscillatory type has the highest rates than the two other types. This is also the case for the propterygium rates. In all these comparisons for swimming types, a significant effect on the shape variables is discernible, except for the metapterygium configuration.

### (e) Morphological convergence

We next sought to investigate a possible pattern of evolutionary convergence among Torpediniformes, Rhinopristiformes, sclerorhynchids and Jurassic batoids, because of their high overlap in the morphospace. We compared the clustering results of the tanglegram of the Procrustes distances dendrogram and the calibrated phylogeny. We identified a cluster that converges towards the shape of Jurassic Batoid and Rhinopristiformes (Supplementary Figure 1). Another group formed by †*Cyclobatis* plus Myliobatiformes, and a group consisting of sclerorhynchids, Pristidae, Squatiniformes, and Pristiophoriformes. Therefore, we explored convergence patterns among these groups. Overall, the results indicate that there is no convergence in the shape trajectories between sharks and batoids when comparing both clades’ nodes (Table 1). We only found evidence of convergence when comparing the node of †*Cyclobatis*, which converges to the node of †*Asterotrygon* and †*Heliobatis*. Moreover, the results from convergence using the cluster groups suggest convergent trajectories between extant Rhinopristiformes and Jurassic batoids, and between extant and extinct guitarfish-like species. We found no convergence between extinct Rhinopristiformes and Jurassic batoids (Table 1). Likewise, convergence was found between members of Sclerorhynchidae, Pristidae, and Pristiophoriformes, and between Squatiniformes and Sclerorhynchidae (Table 1).

**Table 1.** Results from the convergence tests on the state condition comparing groups (State1 and State2) and indicating the significance of the shared angle trajectories between the compared groups. Significant results are in bold at a < 0.05 threshold.

| State 1 | State 2 | Angle state | Angle state time | p angle state | p angle state time |
| --- | --- | --- | --- | --- | --- |
| E_Guit | F_Guit | 27.438 | 0.188 | 0.002 | **0.016** |
| E_Guit | Jbatoid | 60.719 | 0.224 | 0.02 | **0.043** |
| F_Guit | Jbatoid | 49.452 | 0.276 | 0.018 | 0.332 |
| Pristidae | Scle | 43.406 | 0.164 | 0.017 | **0.021** |
| Pristidae | Ppho | 58.099 | 0.103 | 0.055 | **0.002** |
| Scle | Ppho | 30.881 | 0.065 | 0.006 | **0.001** |
| Scle | Ange | 30.969 | 0.073 | 0.001 | **0.001** |

### (f) Modularity and Integration

The modularity analyses suggest that there is strong support for a separation of the elements as individual modules, i.e., each skeletal element covaries independently from each other (Supplementary Table 11). Although the configuration considering the propterygium and metapterygium as a single module shows some support (Supplementary Table 11), the effect size indicates that the separation of each element is better supported. By comparing the configuration between the groups of batoids, we found that both members of Myliobatiformes and Rajiformes present a higher modularity signal compared to those of Rhinopristiformes and Torpediniformes (Table 2). When we compared the modularity signal of the extant batoids, we found that members of both Rajiformes and Rhinopristiformes appear to display a higher signal than the other two orders, and the signal is significant (Table 2). The comparison between extant and extinct batoids suggests that fossil taxa have a higher modularity signal, although the effect is not significant (Table 2). When comparing batoids and dorsoventrally flattened sharks, we found that the batoids have a higher signal with a significant effect. The integration strength signal for the groups described above, shows that among extant batoid orders, members of Rajiformes and Myliobatiformes are more integrated than the other two extant orders (Table 3). Extant batoids appear to display a much higher integration than fossil forms. Finally, the comparison between batoids and dorsoventrally flattened sharks indicates that batoids present a more integrated phenotype in the pectoral fin than sharks (Table 3). All these results are consistent with the findings from the global integration test for all comparisons we made (Table 3).

**Table 2.**
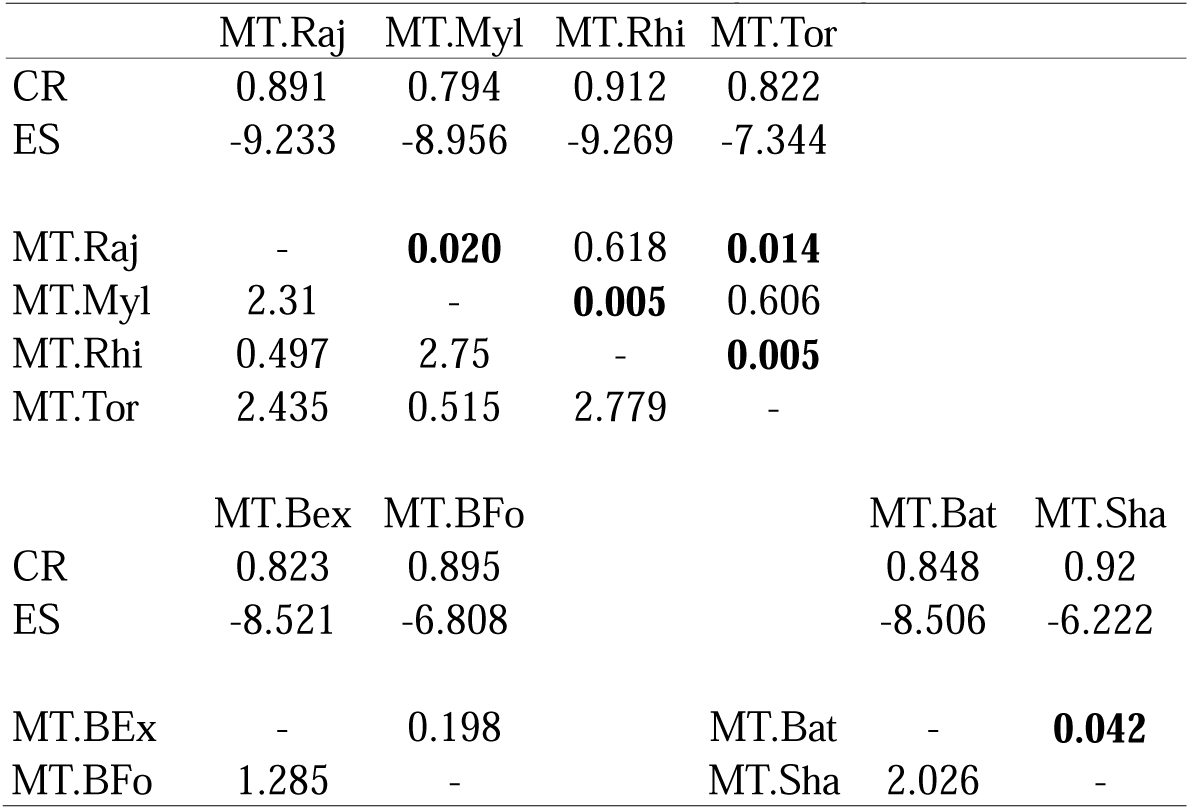
Results from the modularity tests between each group. CR = covariance ratio. ES = Effect size. The effect size is indicated in the lower triangle of each paired comparison and p values on the upper half of the triangle. Significant results are in bold at a < 0.05 threshold.

**Table 3.** Results from the integration test between each group. ES = Effect size. P-pls = r partial least squares value, GI = global integration value. The effect size is indicated in the lower triangle of each paired comparison and p values on the upper half of the triangle. Significant results are in bold at a < 0.05 threshold.

|  | IT.Raj | IT.Myl | IT.Tor | IT.Rhi |  |  |
| --- | --- | --- | --- | --- | --- | --- |
| ES | 5.717 | 5.696 | 3.224 | 4.37 |  |  |
| R-pls | 0.903 | 0.862 | 0.88 | 0.951 |  |  |
| GI | -0.998 | -1.103 | -1.061 | -1.015 |  |  |
| IT.Raj | - | 0.686 | **0.002** | 0.248 |  |  |
| IT.Myl | 0.403 | - | **0.005** | 0.427 |  |  |
| IT.Tor | 3.033 | 2.751 | - | 0.076 |  |  |
| IT.Rhi | 1.153 | 0.794 | 1.771 | - |  |  |
|  | IT.BEx | IT.BFo |  |  | IT.Bat | IT.Sha |
| ES | 7.581 | 3.524 |  |  | 6.771 | 2.678 |
| R-pls | 0.899 | 0.932 |  |  | 0.917 | 0.947 |
| GI | -1.182 | -1.016 |  |  | -1.154 | -0.915 |
| IT.BEx | - | **0.00002** |  | IT.Bat | - | **0.00002** |
| IT.BFo | 4.185 | - |  | IT.Sha | 4.2 | - |

## 4. Discussion

A fundamental aspect of evolutionary biology is to understand the drivers that have led to the evolution of novel phenotypes. As a mean for dispersion, the fins in aquatic organisms represent a structure that might be under several selective pressures and thus responding differently in each individual clade resulting in convergences (69, 70). This results in differently adopted strategies, which ultimately have an impact on their patterns of morphological disparity and evolution. Batoids display a wide array of habitat distribution and swimming types, from the deep-sea/walking skates, to the pelagic/”flying-like” manta rays. It is necessary to understand the impact of such diverse strategies and habitats, as well as processes like modularity and integration in deep time, shaping the evolutionary history of their pectoral fin. We approached this by sampling all possible modern and fossil species encompassing all possible habitats and swimming types. The pectoral fins, specifically the basal elements, represent a trait with a primary function in movement through the environment. Because of the modular nature of its components, it can inform about specific constraints which have an impact on their disparity and convergence patterns. From our results we found that the pectoral fin among batoids has undergone specific changes for each clade and basal element throughout their history. Despite being the most speciose groups, Rajiformes and Myliobatiformes have lower disparities and in the case of Myliobatiformes also evolutionary rates.

Previous findings from the fin outline shape suggest that members of Myliobatiformes have attained higher disparity (6, 7, 71). We found that this pattern is not necessarily the same for the basal fin elements. This could indicate that the higher integration also observed in such groups can allow other parts like the radials to vary independently and achieve higher disparity. From their appearance in the fossil record, the disparity of basal elements of the pectoral fin in batoids constantly increased. Novel phenotypes found during the Eocene from Monte Bolca (17, 72), for instance, demonstrate that batoids experienced a delayed recovery after the KPg massive extinction. This is also matched by observations of the diversity through time of fossils record (19, 73). Among sharks, recent research shows that high disparity was achieved sometime in the Late Cretaceous (74), which is coincident with our findings of disparity through time. The evolution of novel phenotypes resulting in an increased diversity could be associated with the increase of continental fragmentation observed during the Jurassic and Cretaceous, eventually leading to more nearshore environments (73). This could have led groups like Rajiformes, considered for a long time as an invariant group (75), to diverge towards different phenotypes, such as, for instance †*Cyclobatis* with its morphology resembling that seen among freshwater Myliobatiformes (76). Other contemporary representatives of the group display a more “rajiform shape” (e.g., †*Raja davisi*). These fossils illustrate a wider range of disparity in the past, compared to modern groups. In the case of Rajiformes, these changes can also be seen in their present distribution with only a few members in tropical or subtropical waters, but mostly distributed in either temperate to boreal zones or deep sea (77). On the other hand, a consistent pattern of the body shape is discernable among the sharks analyzed here such as Squatiniformes since their origin in the fossil record (21).

We found that the early diverging members of each main order tend to display a generalized morphology, similar to that seen among guitarfishes. This is supported by the convergence tests we conducted. A convergent trajectory between sharks and batoids was not found, only between sharks and the extinct group of sclerorhynchids. Recent developmental studies suggest that the anterior expansion of the pectoral fin is under the control of the *Shh* signaling pathway (78) and that these changes are associated with chromosome architecture during development (79). In addition, hox expression patterns observed in other groups like bamboo sharks, show that the retinoic acid signaling contributes to the posteriorisation of the pectoral fin (80). This signaling alteration leads to a shark-like fin phenotype in skates (81). Nevertheless, the pectoral fin diversity we show suggests that the modification of the pectoral fin also involves the expansion of single elements like the propterygium and metapterygium. This expansion is lacking among sharks, although up to date no developmental studies have been conducted on either saw sharks or angel sharks to confirm such an interpretation. However, the description of developmental sequences in these groups suggests that fin expansion occurs earlier among batoids than in angel sharks (82 83).

From our analyses, the strength of the phylogenetic signal observed differs when different parts of the pectoral fin were analysed, as well as the evolutionary rates for each structure. As in other vertebrates with highly divergent morphologies (84, 85, 86, 87, 88), changes in batoids may follow a mosaic evolution. In addition to the pectoral fins, other features that are unique for the group like the fusion of the most anterior vertebrae (synarcual), the uniquely found anteorbital cartilage, and the dorsally fused suprascapulae, suggest that several changes occurred during the evolution of the group that led to the present morphologies. Some of the Cretaceous and Eocene taxa belonging to extant groups display morphologies with a mixture of characters between different orders (72). More recently, the so-called aquilopelagic phenotype has been shown to have had evolved together with many other modifications for durophagy (71). The radial elements of the pectoral fin also evolved differently than the basal supporting skeletal elements of the fin depending on the swimming type of each species (60, 89). Nevertheless, some genera like *Gymnura* show that the swimming speed is relevant for displaying a specific swimming type (90). Our results indicate that higher evolutionary rates in the coracoid bar shape present a trend for the groups that display oscillatory swimming type.

It has been suggested that extreme morphologies are achieved due to a higher phenotypic integration, which also can be related to low disparity (38, 39, 41). Our results from the integration test indicate that members of both Rajiformes and Myliobatiformes present the highest integration signal among extant batoids. Interestingly, extant batoids appear to have a higher integration than extinct ones. This pattern also was observed in other structures like the skull in angel sharks (91), where extinct forms tend to display lower levels of integration and higher morphological disparity. We found that extinct sclerorhynchids, which attained extreme morphologies resembling the ones seen among sharks (92, 93) display higher disparity than some modern batoids. By comparing batoids and sharks, the pectoral fin of flattened sharks displays a less integrated phenotype and higher modularity. This would suggest that the processes involved in the evolution of the dorsoventral flattening of sharks might not be similar to the ones experienced by batoids. The results of modularity and integration suggest that both signals are significant for the examined groups. However, it has been shown that both processes are not mutually exclusive and the high integration of one module can allow to promote changes across the entire module (94).

## 4. Conclusion

We have shown that the evolution of the pectoral fin skeleton in batoids, the largest group of cartilaginous fishes, appears to be the result of a mosaic evolution. The recurrent guitarfish-like morphology seen from the Jurassic onwards appears to be an intermittent pattern that has evolved several times in their history. This probably has obscured their phylogenetic relationships in past studies. Here, we demonstrate that these morphologies are the result of convergent evolution into a general shape present in most of the groups. The basal fin skeleton follows a different path than the radial portion of the fin, in this sense, the skeletal anatomy seems to carry a higher phylogenetic signal which can support previous descriptions of fossil forms (95). Altogether, we found that the extreme morphologies and reduced disparity of some of the groups can be explained by a process of phenotypic integration, from which the external morphology can develop into different shapes and environments.

## Acknowledgments

Radiographs and CT data obtained from several museum collections mentioned in Materials section.

## Author contributions

Conceptualization: FAL-R, EV-S, and EM

Data Acquisition: FAL-R, JT, FB, RD, SS

Investigation and Analysis: FAL-R, EV-S, FB

Writing and editing: FAL-R, EV-S and EM

Funding acquisition: FAL-R, EM and JK.

## Data availability and Footnotes

Electronic supplementary material is available online at https://github.com/Faviel-LR/Batoid_Fins

## Cited Literatur

[1] Weigmann, S. (2016). Annotated checklist of the living sharks, batoids and chimaeras (Chondrichthyes) of the world, with a focus on biogeographical diversity. Journal of Fish Biology, 88(3), 837–1037.

[2] Fricke, R., Eschmeyer, W. N., & Van der Laan, R. (2024). Catalog of fishes: genera, species, references. California Academy of Sciences, San Francisco, CA, USA http://researcharchive.calacademy.org/research/ichthyology/catalog/fishcatmain.asp.

[3] Compagno, L. J. (1990). Alternative life-history styles of cartilaginous fishes in time and space. Environmental Biology of Fishes, 28, 33–75.

[4] Last, P. R., de Carvalho, M. R., Corrigan, S., Naylor, G. J., Séret, B., & Yang, L. (2016). The Rays of the World project–an explanation of nomenclatural decisions. Rays of the World. Supplementary information, 1–10.

[5] Da Silva, J. P. C., Vaz, D. F., & De Carvalho, M. R. (2018). Phylogenetic inferences on the systematics of squaliform sharks based on elasmobranch scapular morphology (Chondrichthyes: Elasmobranchii). Zoological Journal of the Linnean Society, 182(3), 614–630.

[6] Martinez, C. M., Rohlf, F. J., & Frisk, M. G. (2016). Re evaluation of batoid pectoral morphology reveals novel patterns of diversity among major lineages. Journal of Morphology, 277(4), 482–493.

[7] Franklin, O., Palmer, C., & Dyke, G. (2014). Pectoral fin morphology of batoid fishes (Chondrichthyes: Batoidea): explaining phylogenetic variation with geometric morphometrics. Journal of Morphology, 275(10), 1173–1186.

[8] Schaefer, J. T., & Summers, A. P. (2005). Batoid wing skeletal structure: novel morphologies, mechanical implications, and phylogenetic patterns. Journal of Morphology, 264(3), 298–313.

[9] Türtscher, J., Jambura, P. L., Villalobos Segura, E., López Romero, F. A., Underwood, C. J., Thies, D., … & Kriwet, J. (2024). Rostral and body shape analyses reveal cryptic diversity of Late Jurassic batomorphs (Chondrichthyes, Elasmobranchii) from Europe. Papers in Palaeontology, 10(2), e1552.

[10] Thiollière, V. (1852). Troisième notice sur les gisements à poissons fossiles situés dans le Jura du département de l’Ain. In Annales des Sciences physiques et naturelles, Lyon (Vol. 2, No. 4, pp. 354–446).

[11] Underwood, C. J. (2006). Diversification of the Neoselachii (Chondrichthyes) during the Jurassic and Cretaceous. Paleobiology, 32(2), 215–235.

[12] Stumpf, S., & Kriwet, J. (2019). A new Pliensbachian elasmobranch (Vertebrata, Chondrichthyes) assemblage from Europe, and its contribution to the understanding of late Early Jurassic elasmobranch diversity and distributional patterns. PalZ, 93(4), 637–658.

[13] Villalobos-Segura, E., & Underwood, C. J. (2020). Radiation and divergence times of Batoidea. Journal of Vertebrate Paleontology, 40(3), e1777147.

[14] Cappetta, H. (1980). Les sélaciens du Crétacé supérieur du Liban. II: Batoïdes.

[15] Kachacha, G., Cuny, G., Azar, D., and Abi Saad, P. 2017. Revision of the fossil batomorphs from the Cretaceous of Lebanon, and their impact on our understanding of the early step of the evolution of the clade. Research and Knowledge, 3(2), 33–37

[16] Marramà, G., Claeson, K. M., Carnevale, G., & Kriwet, J. (2018). Revision of Eocene electric rays (Torpediniformes, Batomorphii) from the Bolca Konservat-Lagerstätte, Italy, reveals the first fossil embryo in situ in marine batoids and provides new insights into the origin of trophic novelties in coral reef fishes. Journal of Systematic Palaeontology, 16(14), 1189–1219.

[17] Marramà, G., Carnevale, G., & Kriwet, J. (2021). Diversity, palaeoecology and palaeoenvironmental significance of the Eocene chondrichthyan assemblages of the Bolca Lagerstätte, Italy. Lethaia, 54(5), 736–751.

[18] Kriwet, J., Kiessling, W., & Klug, S. (2009). Diversification trajectories and evolutionary life-history traits in early sharks and batoids. Proceedings of the Royal Society B: Biological Sciences, 276(1658), 945–951.

[19] Guinot, G., Adnet, S., & Cappetta, H. (2012). An analytical approach for estimating fossil record and diversification events in sharks, skates and rays. PLoS One, 7, e44632.

[20] Lund, R. (1988). New information on *Squatinactis caudispinatus* (Chondrichthyes, Cladodontida) from the Chesterian Bear Gulch Limestone of Montana. Journal of Vertebrate Paleontology, 340–342.

[21] Carvalho, M. D., Kriwet, J., & Thies, D. (2008). A systematic and anatomical revision of Late Jurassic angelsharks (Chondrichthyes: Squatinidae). In Mesozoic Fishes: Systematics and Paleoecology; Arratia, G., Viohl, G., Eds.; Verlag Dr Friedrich Pfeil: Munich, Germany, 4, 469–502.

[22] Egeberg, C. A., Kempster, R. M., Theiss, S. M., Hart, N. S., & Collin, S. P. (2014). The distribution and abundance of electrosensory pores in two benthic sharks: a comparison of the wobbegong shark, *Orectolobus maculatus*, and the angel shark, *Squatina australis*. Marine and Freshwater Research, 65(11), 1003–1008.

[23] Duffin, C. J., Garassino, A., & Pasini, G. (2023). *Squaloraja* Riley 1833 (Holocephala: Squalorajidae) from the Lower Jurassic of Osteno Konservat-Lagerstätte (Como, NW Italy). Natural History Sciences, 10(1).

[24] Renz, A. J., Meyer, A., & Kuraku, S. (2013). Revealing less derived nature of cartilaginous fish genomes with their evolutionary time scale inferred with nuclear genes. PLoS One, 8(6), e66400.

[25] Aschliman, N. C., Nishida, M., Miya, M., Inoue, J. G., Rosana, K. M., & Naylor, G. J. (2012). Body plan convergence in the evolution of skates and rays (Chondrichthyes: Batoidea). Molecular Phylogenetics and Evolution, 63(1), 28–42.

[26] Compagno, L. J. (1977). Phyletic relationships of living sharks and rays. American Zoologist, 17(2), 303–322.

[27] Shirai, S. (1992). Phylogenetic relationships of the angel sharks, with comments on elasmobranch phylogeny (Chondrichthyes, Squatinidae). Copeia, 505–518.

[28] Carvalho, M. R. (1996). Higher-level elasmobranch phylogeny, basal squaleans, and paraphyly. Interrelationships of Fishes, Stiassny, M.L.J., Parenti, L.R., Johnson, G.D., Eds.; Academic Press: London, UK, 1996; pp. 35–62

[29] Douady, C. J., Dosay, M., Shivji, M. S., & Stanhope, M. J. (2003). Molecular phylogenetic evidence refuting the hypothesis of Batoidea (rays and skates) as derived sharks. Molecular Phylogenetics and Evolution, 26(2), 215–221.

[30] Naylor, G. J., Caira, J. N., Jensen, K., Rosana, K. A., Straube, N., & Lakner, C. (2012). Elasmobranch phylogeny: a mitochondrial estimate based on 595 species. Biology of Sharks and Their Relatives, 2, 31–56.

[31] Naylor GJP, Yang L, Corrigan S, Carvalho MR (2016) Phylogeny and classification of rays. In: Last PR, White WT, Carvalho MR, Séret B, Stehmann MFW, Naylor GJP (eds) Rays of the World. CSIRO Publishing, Clayton South, pp 10–15

[32] Amaral, C. R., Pereira, F., Silva, D. A., Amorim, A., & de Carvalho, E. F. (2018). The mitogenomic phylogeny of the Elasmobranchii (Chondrichthyes). Mitochondrial DNA Part a, 29(6), 867–878.

[33] Stein, R. W., Mull, C. G., Kuhn, T. S., Aschliman, N. C., Davidson, L. N., Joy, J. B., … & Mooers, A. O. (2018). Global priorities for conserving the evolutionary history of sharks, rays and chimaeras. Nature Ecology & Evolution, 2(2), 288–298.

[34] Kousteni, V., Mazzoleni, S., Vasileiadou, K., & Rovatsos, M. (2021). Complete mitochondrial DNA genome of nine species of sharks and rays and their phylogenetic placement among modern elasmobranchs. Genes, 12(3), 324.

[35] Villalobos-Segura, E., Marramà, G., Carnevale, G., Claeson, K. M., Underwood, C. J., Naylor, G. J., & Kriwet, J. (2022). The phylogeny of rays and skates (Chondrichthyes: Elasmobranchii) based on morphological characters revisited. Diversity, 14(6), 456.

[36] Hoffmann, S. L., Buser, T. J., & Porter, M. E. (2020). Comparative morphology of shark pectoral fins. Journal of Morphology, 281(11), 1501–1516.

[37] Klingenberg, C. P. (2014). Studying morphological integration and modularity at multiple levels: concepts and analysis. Philosophical Transactions of the Royal Society B: Biological Sciences, 369(1649), 20130249.

[38] Goswami, A., Smaers, J. B., Soligo, C., & Polly, P. D. (2014). The macroevolutionary consequences of phenotypic integration: from development to deep time. Philosophical Transactions of the Royal Society B: Biological Sciences, 369(1649), 20130254.

[39] Felice, R. N., Randau, M., & Goswami, A. (2018a). A fly in a tube: macroevolutionary expectations for integrated phenotypes. Evolution, 72(12), 2580–2594.

[40] Zelditch, M. L., & Goswami, A. (2021). What does modularity mean? Evolution & Development, 23(5), 377–403.

[41] Guillerme, T., Cooper, N., Brusatte, S. L., Davis, K. E., Jackson, A. L., Gerber, S., … & Donoghue, P. C. (2020). Disparities in the analysis of morphological disparity. Biology Letters, 16(7), 20200199.

[42] Erwin, D. H. (2007). Disparity: morphological pattern and developmental context. Palaeontology, 50(1), 57–73.

[43] Puttick, M. N., Guillerme, T., & Wills, M. A. (2020). The complex effects of mass extinctions on morphological disparity. Evolution, 74(10), 2207–2220.

[44] Wagner, G. P., Pavlicev, M., & Cheverud, J. M. (2007). The road to modularity. Nature Reviews Genetics, 8(12), 921–931.

[45] Gayford, J. H., Brazeau, M. D., & Naylor, G. J. (2024). Evolutionary trends in the elasmobranch neurocranium. Scientific Reports, 14(1), 11471.

[46] Kamminga, P., De Bruin, P. W., Geleijns, J., & Brazeau, M. D. (2017). X-ray computed tomography library of shark anatomy and lower jaw surface models. Scientific Data, 4(1), 1–6.

[47] Froese, R. and D. Pauly. Editors. (2024). FishBase. World Wide Web electronic publication. www.fishbase.org, version (02/2024).

[48] McClennen M, Jenkins J, Uhen M (2024). Paleobiology Database. Version 1.3. Paleobiology Database. Occurrence dataset 10.15468/jfqhiu

[49] Madisson WP, Madisson DR. 2023 Mesquite: a modular system for evolutionary analysis. Version 3.81.

[50] Goloboff PA, Morales ME. 2023 TNT version 1.6, with a graphical interface for MacOS and Linus, including new routines in parallel. Cladistics, 39, 144–153.

[51] Swofford, D. L. PAUP*. 2003 Phylogenetic Analysis Using Parsimony (*and Other Methods) Version 4.0a166 (Sinauer Associates, 2003).

[52] Brazeau, M. D., Giles, S., Dearden, R. P., Jerve, A., Ariunchimeg, Y. A., Zorig, E., … & Castiello, M. (2020). Endochondral bone in an Early Devonian ‘placoderm’ from Mongolia. Nature Ecology & Evolution, 4(11), 1477–1484.

[53] Rohlf, F. J. (2017). Program TpsDig, version 2.31. Department of Ecology and Evolution, State University of New York at Stony Brook, Stony Brook, NY, USA.

[54] Rohlf, F. J., & Slice, D. (1990). Extensions of the Procrustes method for the optimal superimposition of landmarks. Systematic zoology, 39(1), 40–59.

[55] Gunz, P., & Mitteroecker, P. (2013). Semilandmarks: a method for quantifying curves and surfaces. Hystrix, the Italian Journal of Mammalogy, 24(1), 103–109.

[56] Adams, D. C., & Otárola Castillo, E. (2013). geomorph: an R package for the collection and analysis of geometric morphometric shape data. Methods in ecology and evolution, 4(4), 393–399.

[57] Collyer, M. L., & Adams, D. C. (2021). Phylogenetically aligned component analysis. Methods in Ecology and Evolution, 12(2), 359–372.

[58] Guillerme, T. (2018). dispRity: a modular R package for measuring disparity. Methods in Ecology and Evolution, 9(7), 1755–1763.

[59] Martinez, C. M., Friedman, S. T., Corn, K. A., Larouche, O., Price, S. A., & Wainwright, P. C. (2021). The deep sea is a hot spot of fish body shape evolution. Ecology Letters, 24(9), 1788–1799.

[60] Rosenberger, L. J. (2001). Pectoral fin locomotion in batoid fishes: undulation versus oscillation. Journal of Experimental Biology, 204(2), 379–394.

[61] Castiglione, S., Tesone, G., Piccolo, M., Melchionna, M., Mondanaro, A., Serio, C., … & Raia, P. (2018). A new method for testing evolutionary rate variation and shifts in phenotypic evolution. Methods in Ecology and Evolution, 9(4), 974–983.

[62] Louca, S., & Doebeli, M. (2018). Efficient comparative phylogenetics on large trees. Bioinformatics, 34(6), 1053–1055.

[63] Revell, L. J. (2024). phytools 2.0: an updated R ecosystem for phylogenetic comparative methods (and other things). PeerJ, 12, e16505.

[64] Clavel, J., Escarguel, G., & Merceron, G. (2015). mvMORPH: an R package for fitting multivariate evolutionary models to morphometric data. Methods in Ecology and Evolution, 6(11), 1311–1319.

[65] Castiglione, S., Serio, C., Tamagnini, D., Melchionna, M., Mondanaro, A., Di Febbraro, M., … & Raia, P. (2019). A new, fast method to search for morphological convergence with shape data. PloS One, 14(12), e0226949.

[66] Adams, D. C., & Collyer, M. L. (2019). Comparing the strength of modular signal, and evaluating alternative modular hypotheses, using covariance ratio effect sizes with morphometric data. Evolution, 73(12), 2352–2367.

[67] Adams, D. C., & Collyer, M. L. (2016). On the comparison of the strength of morphological integration across morphometric datasets. Evolution, 70(11), 2623–2631.

[68] Bookstein, F. L. (2015). Integration, disintegration, and self-similarity: characterizing the scales of shape variation in landmark data. Evolutionary Biology, 42, 395–426.

[69] Donley, J. M., Sepulveda, C. A., Konstantinidis, P., Gemballa, S., & Shadwick, R. E. (2004). Convergent evolution in mechanical design of lamnid sharks and tunas. Nature, 429(6987), 61–65.

[70] Fish, F. E. (2023). Aquatic locomotion: Environmental constraints that drive convergent evolution. In Convergent evolution: animal form and function (pp. 477–522). Cham: Springer International Publishing.

[71] Marramà, G., Villalobos Segura, E., Zorzin, R., Kriwet, J., & Carnevale, G. (2023). The evolutionary origin of the durophagous pelagic stingray ecomorph. Palaeontology, 66(4), e12669.

[72] Marramà, G., Schultz, O., & Kriwet, J. (2019). A new Miocene skate from the central Paratethys (Upper Austria): the first unambiguous skeletal record for the Rajiformes (Chondrichthyes: Batomorphii). Journal of Systematic Palaeontology, 17(11), 937–960.

[73] Guinot, G., & Cavin, L. (2020). Distinct responses of elasmobranchs and ray-finned fishes to long-term global change. Frontiers in Ecology and Evolution, 7, 513.

[74] Sternes, P. C., Schmitz, L., & Higham, T. E. (2024). The rise of pelagic sharks and adaptive evolution of pectoral fin morphology during the Cretaceous. Current Biology. 34, 1–9

[75] McEachran, J. D., & Dunn, K. A. (1998). Phylogenetic analysis of skates, a morphologically conservative clade of elasmobranchs (Chondrichthyes: Rajidae). Copeia, 271–290.

[76] Forey, P. L., Yi, L., Patterson, C., & Davies, C. E. (2003). Fossil fishes from the Cenomanian (Upper Cretaceous) of Namoura, Lebanon. Journal of Systematic Palaeontology, 1(4), 227.

[77] Ebert, D.A. & Compagno, L.J.V. (2007) Biodiversity and systematics of skates (Chondrichthyes: Rajiformes: Rajoidei). Environmental Biology of Fishes, 80, 111–124.

[78] Dahn, R. D., Davis, M. C., Pappano, W. N., & Shubin, N. H. (2007). Sonic hedgehog function in chondrichthyan fins and the evolution of appendage patterning. Nature, 445(7125), 311–314.

[79] Marlétaz, F., de la Calle-Mustienes, E., Acemel, R. D., Paliou, C., Naranjo, S., Martínez-García, P. M., … & Gómez-Skarmeta, J. L. (2023). The little skate genome and the evolutionary emergence of wing-like fins. Nature, 616(7957), 495–503.

[80] Onimaru, K., Kuraku, S., Takagi, W., Hyodo, S., Sharpe, J., & Tanaka, M. (2015). A shift in anterior–posterior positional information underlies the fin-to-limb evolution. eLife, 4, e07048.

[81] Nakamura, T., Klomp, J., Pieretti, J., Schneider, I., Gehrke, A. R., & Shubin, N. H. (2015). Molecular mechanisms underlying the exceptional adaptations of batoid fins. Proceedings of the National Academy of Sciences, 112(52), 15940–15945.

[82] Natanson, L. J., & Cailliet, G. M. (1986). Reproduction and development of the Pacific angel shark, *Squatina californica*, off Santa Barbara, California. Copeia, 987–994.

[83] Maxwell, E. E., Fröbisch, N. B., & Heppleston, A. C. (2008). Variability and conservation in late chondrichthyan development: ontogeny of the winter skate (*Leucoraja ocellata*). The Anatomical Record, 291(9), 1079–1087.

[84] Felice, R. N., & Goswami, A. (2018b). Developmental origins of mosaic evolution in the avian cranium. Proceedings of the National Academy of Sciences, 115(3), 555–560.

[85] Bardua, C., Fabre, A. C., Bon, M., Das, K., Stanley, E. L., Blackburn, D. C., & Goswami, A. (2020). Evolutionary integration of the frog cranium. Evolution, 74(6), 1200–1215.

[86] Coombs, E. J., Felice, R. N., Clavel, J., Park, T., Bennion, R. F., Churchill, M., … & Goswami, A. (2022). The tempo of cetacean cranial evolution. Current Biology, 32(10), 2233–2247.

[87] Larouche, O., Gartner, S. M., Westneat, M. W., & Evans, K. M. (2023). Mosaic evolution of the skull in labrid fishes involves differences in both tempo and mode of morphological change. Systematic Biology, 72(2), 419–432.

[88] Law, C. J., Hlusko, L. J., & Tseng, Z. J. (2024). Uncovering the mosaic evolution of the carnivoran skeletal system. Biology Letters, 20(1), 20230526.

[89] Hall, K. C., Hundt, P. J., Swenson, J. D., Summers, A. P., & Crow, K. D. (2018). The evolution of underwater flight: the redistribution of pectoral fin rays, in manta rays and their relatives (Myliobatidae). Journal of Morphology, 279(8), 1155–1170.

[90] Kim, H. S., Lee, J. Y., Chu, W. S., & Ahn, S. H. (2017). Design and fabrication of soft morphing ray propulsor: undulator and oscillator. Soft Robotics, 4(1), 49–60.

[91] López-Romero, F. A., Stumpf, S., Pfaff, C., Marramà, G., Johanson, Z., & Kriwet, J. (2020). Evolutionary trends of the conserved neurocranium shape in angel sharks (Squatiniformes, Elasmobranchii). Scientific Reports, 10(1), 12582.

[92] Wueringer, B. E., Squire, L., & Collin, S. P. (2009). The biology of extinct and extant sawfish (Batoidea: Sclerorhynchidae and Pristidae). Reviews in Fish Biology and Fisheries, 19, 445–464.

[93] Villalobos Segura, E., Underwood, C. J., & Ward, D. J. (2021). The first skeletal record of the enigmatic Cretaceous sawfish genus *Ptychotrygon* (Chondrichthyes, Batoidea) from the Turonian of Morocco. Papers in Palaeontology, 7(1), 353–376.

[94] Goswami, A., & Polly, P. D. (2010). The influence of modularity on cranial morphological disparity in Carnivora and Primates (Mammalia). PloS One, 5(3), e9517.

[95] Sansom, R. S., & Wills, M. A. (2013). Fossilization causes organisms to appear erroneously primitive by distorting evolutionary trees. Scientific Reports, 3(1), 2545.

