## Supplementary Material for "Evolution of the Batoidea Pectoral Fin Skeleton: Convergence, Modularity, and Integration Driving Disparity Trends"

López-Romero Faviel A.,[1] Villalobos-Segura Eduardo,[2] Türtscher Julia,[2, 3] Berio Fidji,[4] Stumpf Sebastian,[2] Dearden Richard P.,[5,6] Kriwet Jürgen,[2, 3] and Maldonado Ernesto[1]*.

1. EvoDevo Research Group, Unidad de Sistemas Arrecifales, Instituto de Ciencias del Mar y Limnología, Universidad Nacional Autónoma de México, Puerto Morelos, Quintana Roo, México;

2 University of Vienna, Faculty of Earth Sciences, Geography and Astronomy, Department of Palaeontology, Evolutionary Morphology Research Group, Josef-Holaubek-Platz 2, 1190, Vienna, Austria

3 University of Vienna, Vienna Doctoral School of Ecology and Evolution (VDSEE), Djerassiplatz 1, 1030, Vienna, Austria

4 Department of Zoology, Stockholm University, Svante Arrhenius väg 18B, 114 18, Stockholm, Sweden.

5 Vertebrate Evolution, Development, and Ecology, Naturalis Biodiversity Center, Darwinweg 2, Leiden, 2333 CR, The Netherlands

6 University of Birmingham, School of Geography, Earth & Environmental Sciences, University of Birmingham, Edgbaston, Birmingham B15 2TT, UK

*Correspondence to: Ernesto Maldonado.
Unidad Académica de Sistemas Arrecifales, Instituto de Ciencias del Mar y Limnología,
Universidad Nacional Autónoma de México, C.P. 77580 Puerto Morelos, Quintana Roo, México.
**

**Supplementary Table 1.** Landmark definitions used for the present study.

| Landmark/Curve | |
|---|---|
| L1 | Anterior midline of the coracoid bar |
| C1 | Anterior curve of the coracoid bar |
| L2 | Inner coracoid bar-propterygium joint |
| C2 | Inner curve of the first propterygium segment |
| L3 | Anterior inner propterygium joint with second propterygium segment |
| L4 | Anterior outer propterygium joint with second propterygium segment |
| C3 | Outer curve of the first propterygium segment |
| L5 | Outer coracoid bar-propterygium joint |
| L6 | Posterior midline of the coracoid bar |
| C4 | Posterior curve of the coracoid bar |
| L7 | Inner coracoid bar-metapterygium joint |
| C5 | Inner curve of the first metapterygium segment |
| L8 | Posterior inner metapterygium joint with second metapterygium segment |
| L9 | Posterior outer metapterygium joint with second metapterygium segment |
| C6 | Outer curve of the first metapterygium segment |
| L10 | Outer coracoid bar-metapterygium joint |
| C7 | Coracoid bar - mesopterygium joint curve |

**Supplementary Table 2.** Morphological disparity as sum of variances for all groups. Subsets of taxonomic groups, temporal groups, habitat categories, and swimming categories. All (All landmark configurations), CB (Coracoid Bar configuration), PR (Propterygium), MT (Metapterygium). N (number of samples per group) obs (observed disparity) bs.median (bootstrapped median disparity)

| | All | | | CB | | PR | | MT | |
|---|---|---|---|---|---|---|---|---|---|
| subsets | n | obs | bs.median | obs | bs.median | obs | bs.median | obs | bs.median |
| Rhinop | 38 | 0.018 | 0.018 | 0.01 | 0.01 | 0.014 | 0.014 | 0.006 | 0.006 |
| Myliob | 131 | 0.02 | 0.02 | 0.039 | 0.039 | 0.024 | 0.023 | 0.006 | 0.006 |
| Torped | 35 | 0.036 | 0.035 | 0.012 | 0.011 | 0.032 | 0.03 | 0.041 | 0.04 |
| Rajifo | 91 | 0.012 | 0.012 | 0.026 | 0.026 | 0.015 | 0.015 | 0.004 | 0.004 |
| Orecto | 5 | 0.039 | 0.032 | 0.025 | 0.02 | 0.081 | 0.059 | 0.014 | 0.01 |
| Pristi | 6 | 0.019 | 0.015 | 0.012 | 0.011 | 0.05 | 0.044 | 0.021 | 0.018 |
| Squati | 16 | 0.005 | 0.005 | 0.003 | 0.003 | 0.015 | 0.014 | 0.007 | 0.006 |
| Cr_Raj | 3 | 0.004 | 0.002 | 0.017 | 0.012 | 0.026 | 0.025 | 0.005 | 0.004 |
| Cr_Scl | 6 | 0.025 | 0.02 | 0.013 | 0.011 | 0.024 | 0.021 | 0.035 | 0.031 |
| Cr_Rhi | 12 | 0.02 | 0.019 | 0.037 | 0.03 | 0.013 | 0.012 | 0.007 | 0.007 |
| Eo_Myl | 9 | 0.01 | 0.01 | 0.029 | 0.027 | 0.012 | 0.012 | 0.003 | 0.003 |
| Jr_Squ | 3 | 0.018 | 0.013 | 0.018 | 0.011 | 0.044 | 0.034 | 0.01 | 0.006 |
| Jr_Bat | 11 | 0.011 | 0.01 | 0.017 | 0.015 | 0.009 | 0.008 | 0.005 | 0.005 |
| Batoids | 288 | 0.072 | 0.072 | 0.07 | 0.069 | 0.063 | 0.063 | 0.023 | 0.023 |
| Sharks | 25 | 0.022 | 0.022 | 0.017 | 0.016 | 0.082 | 0.08 | 0.015 | 0.014 |
| Cr_Bat | 21 | 0.055 | 0.054 | 0.049 | 0.047 | 0.028 | 0.028 | 0.023 | 0.02 |
| Eo_Bat | 10 | 0.067 | 0.062 | 0.049 | 0.044 | 0.038 | 0.037 | 0.026 | 0.025 |
| Jr_Sha | 5 | 0.047 | 0.042 | 0.033 | 0.029 | 0.086 | 0.065 | 0.009 | 0.008 |
| Jr_Bat | 11 | 0.011 | 0.01 | 0.017 | 0.015 | 0.009 | 0.008 | 0.005 | 0.005 |
| Deep | 82 | 0.025 | 0.026 | 0.029 | 0.028 | 0.021 | 0.021 | 0.01 | 0.008 |
| FreshW | 19 | 0.006 | 0.006 | 0.011 | 0.01 | 0.009 | 0.009 | 0.003 | 0.002 |
| Reef | 41 | 0.073 | 0.07 | 0.088 | 0.084 | 0.056 | 0.055 | 0.038 | 0.036 |
| Shelf | 146 | 0.071 | 0.071 | 0.087 | 0.086 | 0.051 | 0.049 | 0.027 | 0.026 |
| Undul | 191 | 0.05 | 0.05 | 0.041 | 0.041 | 0.049 | 0.049 | 0.005 | 0.004 |
| Oscil | 36 | 0.008 | 0.008 | 0.022 | 0.021 | 0.012 | 0.012 | 0.009 | 0.009 |
| AxUnd | 61 | 0.059 | 0.058 | 0.018 | 0.018 | 0.034 | 0.034 | 0.051 | 0.049 |

**Supplementary Table 3.** Pairwise comparison of the disparity by groups for the whole landmark configuration. Upper triangle indicates the P value, lower triangle indicates the statistic W of the Wilcoxon test. Bold indicates p<0.05

| | Rhinop | Myliob | Torped | Rajifo | Orecto | Pristi | Squati | Cr_Raj | Cr_Scl | Cr_Rhi | Eo_Myl | Jr_Squ | Jr_Bat |
|---|---|---|---|---|---|---|---|---|---|---|---|---|---|
| Rhinop | | **>0.001** | **>0.001** | **>0.001** | **>0.001** | **0.008** | **>0.001** | **>0.001** | **0.0333** | 1 | **>0.001** | **>0.001** | **>0.001** |
| Myliob | 3007 | | **>0.001** | **>0.001** | **>0.001** | **>0.001** | **>0.001** | **>0.001** | 1 | **0.0012** | **>0.001** | **>0.001** | **>0.001** |
| Torped | 183 | 335 | | **>0.001** | 1 | **>0.001** | **>0.001** | **>0.001** | **>0.001** | **>0.001** | **>0.001** | **>0.001** | **>0.001** |
| Rajifo | 9854 | 10000 | 9992 | | **>0.001** | **>0.001** | **>0.001** | **>0.001** | **>0.001** | **>0.001** | **>0.001** | 0.1271 | **>0.001** |
| Orecto | 732 | 920 | 5938 | 58 | | **>0.001** | **>0.001** | **>0.001** | **>0.001** | **>0.001** | **>0.001** | **>0.001** | **>0.001** |
| Pristi | 6590 | 7701 | 9862 | 2348 | 9491 | | **>0.001** | **>0.001** | **>0.001** | **>0.001** | **>0.001** | **0.01802** | **>0.001** |
| Squati | 10000 | 10000 | 10000 | 10000 | 10000 | 9905 | | **>0.001** | **>0.001** | **>0.001** | **>0.001** | **>0.001** | **>0.001** |
| Cr_Raj | 10000 | 10000 | 10000 | 10000 | 10000 | 10000 | 9770 | | **>0.001** | **>0.001** | **>0.001** | **>0.001** | **>0.001** |
| Cr_Scl | 3558 | 4880 | 9424 | 401 | 8725 | 2550 | 0 | 0 | | 0.169 | **>0.001** | **>0.001** | **>0.001** |
| Cr_Rhi | 4690 | 6764 | 9799 | 119 | 9264 | 3291 | 0 | 0 | 6255 | | **>0.001** | **>0.001** | **>0.001** |
| Eo_Myl | 9780 | 9958 | 9979 | 6990 | 9882 | 8306 | 1246 | 171 | 9611 | 9795 | | **0.0022** | 1 |
| Jr_Squ | 7927 | 9069 | 9936 | 3714 | 9647 | 6503 | 1100 | 1083.5 | 8429 | 8138 | 3292 | | **>0.001** |
| Jr_Bat | 9965 | 10000 | 10000 | 8552 | 9994 | 8693 | 20 | 0 | 9839 | 9977 | 4938 | 6450 | |

| | Batoid | Sharks | Cr_Bat | Eo_Bat | Jr_Sha | Jr_Bat |
|---|---|---|---|---|---|---|
| Batoids | | **>0.001** | **>0.001** | **>0.001** | **>0.001** | **>0.001** |
| Sharks | 10000 | | **>0.001** | **0.0022** | **>0.001** | **>0.001** |
| Cr_Bat | 9683 | 39 | | 0.11734 | **>0.001** | **>0.001** |
| Eo_Bat | 7194 | 3448 | 3911 | | **>0.001** | **>0.001** |
| Jr_Sha | 9990 | 1973 | 7384 | 6541 | | **>0.001** |
| Jr_Bat | 10000 | 9989 | 10000 | 8376 | 9864 | |

| | Deep | Fresh W | Reef | Shelf |
|---|---|---|---|---|
| Deep | | **>0.001** | **>0.001** | **>0.001** |
| FreshW | 10000 | | **>0.001** | **>0.001** |
| Reef | 1 | 0 | | 1 |
| Shelf | 0 | 0 | 4505 | |

| | Undul | Oscil | AxUnd |
|---|---|---|---|
| Undul | | **>0.001** | **>0.001** |
| Oscil | 10000 | | **>0.001** |
| AxUnd | 52 | 0 | |

**Supplementary Table 4.** Pairwise comparison of the disparity by groups for the coracoid bar landmark configuration. Upper triangle indicates the P value, lower triangle indicates the statistic W of the Wilcoxon test. Bold indicates p<0.05

| | Rhinop | Myliob | Torped | Rajifo | Orecto | Pristi | Squati | Cr_Raj | Cr_Scl | Cr_Rhi | Eo_Myl | Jr_Squ | Jr_Bat |
|---|---|---|---|---|---|---|---|---|---|---|---|---|---|
| Rhinop | | **>0.001** | **>0.001** | **>0.001** | **>0.001** | 1 | **>0.001** | 1 | 1 | **>0.001** | **>0.001** | **>0.001** | **>0.001** |
| Myliob | 0 | | **>0.001** | **>0.001** | **>0.001** | **>0.001** | **>0.001** | **>0.001** | **>0.001** | 0.0761 | **>0.001** | **>0.001** | **>0.001** |
| Torped | 2939 | 10000 | | **>0.001** | **>0.001** | 1 | **>0.001** | 1 | 0.1148 | **>0.001** | **>0.001** | 1 | **>0.001** |
| Rajifo | 0 | 9999 | 0 | | **>0.001** | **>0.001** | **>0.001** | **>0.001** | **>0.001** | 0.646 | 1 | **>0.001** | **>0.001** |
| Orecto | 685 | 10000 | 998 | 9215 | | **>0.001** | **>0.001** | **>0.001** | **>0.001** | **>0.001** | **>0.001** | **>0.001** | **>0.001** |
| Pristi | 4323 | 10000 | 5842 | 10000 | 9125 | | **>0.001** | 0.4779 | 1 | **>0.001** | **>0.001** | 0.265 | **>0.001** |
| Squati | 10000 | 10000 | 10000 | 10000 | 10000 | 10000 | | **>0.001** | **>0.001** | **>0.001** | **>0.001** | **>0.001** | **>0.001** |
| Cr_Raj | 4080 | 10000 | 4736 | 10000 | 8692 | 3882 | 600 | | 0.2168 | **>0.001** | **>0.001** | 1 | **>0.001** |
| Cr_Scl | 4885 | 10000 | 6302 | 10000 | 9240 | 5286 | 50 | 6220 | | **>0.001** | **>0.001** | **0.0057** | **>0.001** |
| Cr_Rhi | 589 | 6350 | 921 | 3919 | 2674 | 814 | 0 | 972 | 678 | | 1 | **>0.001** | **>0.001** |
| Eo_Myl | 0 | 9774 | 34 | 4145 | 1438 | 57 | 0 | 174 | 28 | 5810 | | **>0.001** | **>0.001** |
| Jr_Squ | 2898 | 10000 | 4151 | 10000 | 8537 | 3805 | 900 | 4777 | 3382 | 8855 | 9792 | | **>0.001** |
| Jr_Bat | 202 | 10000 | 1178 | 10000 | 7905 | 1259 | 0 | 3104 | 979 | 8410 | 9747 | 2945 | |

| | Batoids | Sharks | Cr_Bat | Eo_Bat | Jr_Sha | Jr_Bat |
|---|---|---|---|---|---|---|
| Batoids | | **>0.001** | **>0.001** | **>0.001** | **>0.001** | **>0.001** |
| Sharks | 10000 | | **>0.001** | **>0.001** | **>0.001** | **0.0078** |
| Cr_Bat | 9908 | 10 | | 1 | **>0.001** | **>0.001** |
| Eo_Bat | 8901 | 150 | 5669 | | **>0.001** | **>0.001** |
| Jr_Sha | 10000 | 3080 | 8989 | 7594 | | **>0.001** |
| Jr_Bat | 10000 | 6420 | 9999 | 9923 | 6990 | |

| | Deep | FreshW | Reef | Shelf |
|---|---|---|---|---|
| Deep | | **>0.001** | **>0.001** | **>0.001** |
| FreshW | 10000 | | **>0.001** | **>0.001** |
| Reef | 0 | 0 | | 5.10E-02 |
| Shelf | 0 | 0 | 3924 | |

| | Undul | Oscil | AxUnd |
|---|---|---|---|
| Undul | | **>0.001** | **>0.001** |
| Oscil | 10000 | | **>0.001** |
| AxUnd | 10000 | 8939 | |

**Supplementary Table 5.** Pairwise comparison of the disparity by groups for the propterygium landmark configuration. Upper triangle indicates the P value, lower triangle indicates the statistic W of the Wilcoxon test. Bold indicates p<0.05

| | Rhinop | Myliob | Torped | Rajifo | Orecto | Pristi | Squati | Cr_Raj | Cr_Scl | Cr_Rhi | Eo_Myl | Jr_Squ | Jr_Bat |
|---|---|---|---|---|---|---|---|---|---|---|---|---|---|
| Rhinop | | **>0.001** | **>0.001** | **>0.001** | **>0.001** | **>0.001** | 1 | **0.0185** | **>0.001** | **>0.001** | **>0.001** | **>0.001** | **>0.001** |
| Myliob | 1 | | **>0.001** | **>0.001** | **>0.001** | **>0.001** | **>0.001** | 1 | **>0.001** | **>0.001** | **>0.001** | **>0.001** | **>0.001** |
| Torped | 3 | 1844 | | **>0.001** | **>0.001** | 0.1451 | **>0.001** | **>0.001** | **>0.001** | **>0.001** | **>0.001** | 0.0531 | **>0.001** |
| Rajifo | 2391 | 9976 | 9973 | | **>0.001** | **>0.001** | **>0.001** | **0.0185** | **>0.001** | **>0.001** | **>0.001** | **>0.001** | **>0.001** |
| Orecto | 127 | 309 | 605 | 185 | | **>0.001** | **>0.001** | **>0.001** | **>0.001** | **>0.001** | **>0.001** | **>0.001** | **>0.001** |
| Pristi | 161 | 3052 | 3726 | 390 | 7692 | | **>0.001** | **>0.001** | **>0.001** | **>0.001** | **>0.001** | **0.0064** | **>0.001** |
| Squati | 5111 | 9986 | 9985 | 7550 | 9876 | 9793 | | **0.0185** | **>0.001** | **>0.001** | **>0.001** | **>0.001** | **>0.001** |
| Cr_Raj | 3500 | 5980 | 8659 | 3500 | 9805 | 7601 | 3500 | | 1 | **0.0185** | **0.0185** | **>0.001** | **0.0185** |
| Cr_Scl | 1628 | 7022 | 8926 | 2244 | 9754 | 7976 | 1591 | 5500 | | **>0.001** | **>0.001** | **>0.001** | **>0.001** |
| Cr_Rhi | 7563 | 10000 | 10000 | 9045 | 9936 | 9938 | 7259 | 6500 | 9032 | | 1 | **>0.001** | **>0.001** |
| Eo_Myl | 7863 | 10000 | 10000 | 9214 | 9947 | 9947 | 7609 | 6500 | 9143 | 5611 | | **>0.001** | **>0.001** |
| Jr_Squ | 2900 | 2924 | 3614 | 2900 | 8774 | 6606 | 2831 | 2068 | 2877 | 2739 | 2440 | | **>0.001** |
| Jr_Bat | 9964 | 10000 | 10000 | 10000 | 10000 | 10000 | 9752 | 6500 | 9853 | 9085 | 8076 | 8365 | |

| | Batoids | Sharks | Cr_Bat | Eo_Bat | Jr_Sha | Jr_Bat |
|---|---|---|---|---|---|---|
| Batoids | | **>0.001** | **>0.001** | **>0.001** | 1 | **>0.001** |
| Sharks | 2069 | | **>0.001** | **>0.001** | **0.0031** | **>0.001** |
| Cr_Bat | 10000 | 9949 | | **0.031** | **>0.001** | **>0.001** |
| Eo_Bat | 9073 | 9320 | 3739 | | **>0.001** | **>0.001** |
| Jr_Sha | 4275 | 6516 | 459 | 1547 | | **>0.001** |
| Jr_Bat | 10000 | 10000 | 10000 | 9401 | 10000 | |

| | Deep | Fresh W | Reef | Shelf |
|---|---|---|---|---|
| Deep | | **>0.001** | **>0.001** | **>0.001** |
| FreshW | 10000 | | **>0.001** | **>0.001** |
| Reef | 0 | 0 | | **>0.001** |
| Shelf | 0 | 0 | 7460 | |

| | Undul | Oscil | AxUnd |
|---|---|---|---|
| Undul | | **>0.001** | **>0.001** |
| Oscil | 10000 | | **>0.001** |
| AxUnd | 10000 | 0 | |

**Supplementary Table 6.** Pairwise comparison of the disparity by groups for the metapterygium landmark configuration. Upper triangle indicates the P value, lower triangle indicates the statistic W of the Wilcoxon test. Bold indicates $p<0.05$

| | Rhinop | Myliob | Torped | Rajifo | Orecto | Pristi | Squati | Cr_Raj | Cr_Scl | Cr_Rhi | Eo_Myl | Jr_Squ | Jr_Bat |
|---|---|---|---|---|---|---|---|---|---|---|---|---|---|
| Rhinop | | 1 | **>0.001** | **>0.001** | **>0.001** | **>0.001** | 1 | **>0.001** | **>0.001** | **>0.001** | **>0.001** | 1 | **0.0035** |
| Myliob | 4985 | | **>0.001** | **>0.001** | **>0.001** | **>0.001** | 0.9532 | **>0.001** | **>0.001** | **>0.001** | **>0.001** | 1 | **>0.001** |
| Torped | 0 | 0 | | **>0.001** | **>0.001** | **>0.001** | **>0.001** | **>0.001** | **>0.001** | **>0.001** | **>0.001** | **>0.001** | **>0.001** |
| Rajifo | 9847 | 10000 | 10000 | | **>0.001** | **>0.001** | **>0.001** | **0.0084** | **>0.001** | **>0.001** | **>0.001** | **>0.001** | **>0.001** |
| Orecto | 1811 | 1852 | 10000 | 257 | | **>0.001** | **>0.001** | **>0.001** | **>0.001** | **>0.001** | **>0.001** | **>0.001** | **>0.001** |
| Pristi | 462 | 413 | 9988 | 300 | 2485 | | **>0.001** | **>0.001** | **>0.001** | **>0.001** | **>0.001** | **>0.001** | **>0.001** |
| Squati | 4124 | 3974 | 10000 | 201 | 7930 | 9438 | | **>0.001** | **>0.001** | 0.7406 | **>0.001** | 1 | **>0.001** |
| Cr_Raj | 8758 | 9514 | 10000 | 3420 | 9144 | 9802 | 8868 | | **>0.001** | **>0.001** | **0.00689** | **>0.001** | **>0.001** |
| Cr_Scl | 333 | 345 | 7916 | 0 | 561 | 1477 | 396 | 76 | | **>0.001** | **>0.001** | **>0.001** | **>0.001** |
| Cr_Rhi | 2982 | 2683 | 10000 | 218 | 7621 | 9325 | 3938 | 564 | 9535 | | **>0.001** | 1 | **>0.001** |
| Eo_Myl | 9990 | 10000 | 10000 | 9368 | 9889 | 9708 | 9988 | 6600 | 10000 | 9979 | | **>0.001** | **>0.001** |
| Jr_Squ | 4612 | 4928 | 10000 | 1228 | 7624 | 9185 | 4792 | 2404 | 9672 | 5105 | 1200 | | 0.09 |
| Jr_Bat | 6668 | 6888 | 10000 | 1033 | 8647 | 9621 | 7264 | 2652 | 9784 | 8112 | 255 | 6320 | |

| | Batoid | Sharks | Cr_Bat | Eo_Bat | Jr_Sha | Jr_Bat |
|---|---|---|---|---|---|---|
| Batoids | | **>0.001** | **0.0064** | 1 | **>0.001** | **>0.001** |
| Sharks | 9903 | | **>0.001** | 0.2189 | **>0.001** | **>0.001** |
| Cr_Bat | 6442 | 2232 | | 1 | **>0.001** | **>0.001** |
| Eo_Bat | 4598 | 4000 | 4853 | | 0.0735 | 0.15 |
| Jr_Sha | 10000 | 9875 | 9663 | 6152 | | **>0.001** |
| Jr_Bat | 10000 | 10000 | 9971 | 6055 | 7599 | |

| | Deep | FreshW | Reef | Shelf |
|---|---|---|---|---|
| Deep | | **>0.001** | **>0.001** | **>0.001** |
| FreshW | 9976 | | **>0.001** | **>0.001** |
| Reef | 1 | 0 | | **>0.001** |
| Shelf | 18 | 0 | 9003 | |

| | Undul | Oscil | AxUnd |
|---|---|---|---|
| Undul | | **>0.001** | **>0.001** |
| Oscil | 0 | | **>0.001** |
| AxUnd | 0 | 0 | |

**Supplementary Table 7.** Evolutionary rates and shift by clades, p value indicates the positive (p>0.975) or negative (p<0.025) shifts from the search.shift function in RRphylo. All the configurations and isolated elements comparisons. * Result found with the “auto” parameter in search.shift.

| | Node | Group | rate.difference | p.value |
|---|---|---|---|---|
| Full configuration | | All | 0.001665675 | 0.915 |
| | 406 | Sela | 0.0002815367 | 0.563 |
| | 213 | Bats | 0.0017804218 | 0.94 |
| | 403 | Jbat | 0.0003566254 | 0.892 |
| | 401 | Scle | 0.0014195306 | **0.9995** |
| | 216 | Rhin | 0.002341807 | 0.948 |
| | 384 | Torp | 0.0006783165 | 0.891 |
| | 322 | Raji | 0.0027842153 | **0.999** |
| | 249 | Mylio | 0.001021813 | 0.791 |
| | 249 | Mylio* | -0.0009675225 | **0.001** |
| Coracoid bar | | All | 0.003401622 | 0.995 |
| | 406 | Sela | 0.0004246853 | 0.774 |
| | 213 | Bats | 0.0036484149 | **0.999** |
| | 403 | Jbat | 0.0007212336 | **1** |
| | 401 | Scle | 0.0014043769 | **0.9995** |
| | 216 | Rhin | 0.0055538655 | **1** |
| | 384 | Torp | 0.0004796465 | 0.946 |
| | 322 | Raji | 0.003950205 | **1** |
| | 249 | Mylio | 0.0036225049 | **0.999** |
| Propterygium | | All | 0.002695441 | 0.981 |
| | 406 | Sela | 0.0005301853 | 0.719 |
| | 213 | Bats | 0.0028749438 | **0.987** |
| | 403 | Jbat | 0.0002726882 | 0.806 |
| | 401 | Scle | 0.0022005471 | **0.9865** |
| | 216 | Rhin | 0.002035749 | **0.991** |
| | 384 | Torp | 0.0022996407 | **0.989** |
| | 322 | Raji | 0.0037849555 | **1** |
| | 249 | Mylio | 0.0028783547 | 0.984 |
| Metapterygium | | All | 0.001661485 | 0.997 |
| | 406 | Sela | 0.0004578208 | 0.922 |
| | 213 | Bats | 0.0017612708 | **0.995** |
| | 403 | Jbat | 0.0007751719 | **0.998** |
| | 401 | Scle | 0.0020142031 | **1** |
| | 216 | Rhin | 0.0021474323 | **1** |
| | 384 | Torp | 0.0024454247 | **1** |
| | 322 | Raji | 0.0014385677 | **0.99** |
| | 249 | Mylio | 0.0018033827 | **1** |

**Supplementary table 8.** Results from the overFit function to account for phylogenetic uncertainty. 100 random trees were used. p.shift+ or p.shift- indicates the percentage of times the same shift was found in the random sample. And tested.trees the percentage of trees used. Bold indicates the same result as the one obtained with a single tree.

| | | p.shift+ | p.shift- | tested.trees |
|---|---|---|---|---|
| Full configuration | all.cla | 0 | 0 | 1 |
| | 406 | 0 | 0 | 1 |
| | 213 | 0 | 0 | 1 |
| | 403 | 0 | 0 | 1 |
| | 401 | **1** | 0 | 1 |
| | 216 | 0 | 0 | 1 |
| | 384 | 0 | 0 | 1 |
| | 322 | **1** | 0 | 1 |
| | 249 | 0 | 0 | 1 |
| Coracoid bar | all.cla | 1 | 0 | 1 |
| | 406 | 0 | 0 | 1 |
| | 213 | **1** | 0 | 1 |
| | 403 | **1** | 0 | 1 |
| | 401 | **1** | 0 | 1 |
| | 216 | **0.99** | 0 | 1 |
| | 384 | 0.13 | 0 | 1 |
| | 322 | **1** | 0 | 1 |
| | 249 | **1** | 0 | 1 |
| Propterygium | all.cla | 0.17 | 0 | 1 |
| | 406 | 0 | 0 | 1 |
| | 213 | **0.42** | 0 | 1 |
| | 403 | 0 | 0 | 1 |
| | 401 | **1** | 0 | 1 |
| | 216 | 0.12 | 0 | 1 |
| | 384 | **0.86** | 0 | 1 |
| | 322 | **1** | 0 | 1 |
| | 249 | 0.18 | 0 | 1 |
| Metapterygium | all.clades | 0.96 | 0 | 1 |
| | 406 | 0 | 0 | 1 |
| | 213 | **0.98** | 0 | 1 |
| | 403 | **1** | 0 | 1 |
| | 401 | **1** | 0 | 1 |
| | 216 | **0.99** | 0 | 1 |
| | 384 | **1** | 0 | 1 |
| | 322 | 0.79 | 0 | 1 |
| | 249 | **0.98** | 0 | 1 |

**Supplementary Table 9.** Evolutionary rates as sigma^2 from the Extant set comparing habitat groups and swimming categories for the whole configuration (All), coracoid bar (CB), propterygium (PR) or metapterygium (MT).

| | All | CB | PR | MT |
|---|---|---|---|---|
| Deep | 1.41E-05 | 2.88E-05 | 1.90E-05 | 1.27E-06 |
| Freshwater | 2.08E-05 | 6.86E-05 | 4.97E-05 | 1.61E-06 |
| Reef | 7.68E-05 | 8.41E-05 | 7.13E-05 | 2.20E-05 |
| Shelf | 3.41E-05 | 5.97E-05 | 2.88E-05 | 1.59E-05 |
| AxUnd | 6.11E-05 | 9.37E-06 | 3.40E-05 | 3.74E-05 |
| Oscil | 1.92E-05 | 3.22E-04 | 4.80E-05 | 1.21E-05 |
| Undul | 2.82E-05 | 8.09E-05 | 3.48E-05 | 2.61E-06 |

**Supplementary Table 10.** Results from the manova.gls test for the shape variables and the groups. Bold indicates a significant interaction at p<0.05

| Structure | Group | Test stat | Pr(>Stat) |
|---|---|---|---|
| All | Amb | 0.1245 | 0.29 |
| | Mov | 0.3271 | **0.001** |
| CB | Amb | 0.1715 | **0.001** |
| | Mov | 0.4153 | **0.001** |
| PR | Amb | 0.08498 | **0.041** |
| | Mov | 0.1966 | **0.007** |
| MT | Amb | 0.09454 | 0.26 |
| | Mov | 0.109 | **0.041** |

**Supplementary Table 11.** Effect size pairwise comparison between the modularity hypotheses tested. The CR value is indicated for each configuration. Lower triangle indicates the effect size, and upper triangle the p value. Bold indicates significant ($p<0.05$)

| Sharks and Batoids | No_Modules | MT.AL | MT.GP | MT.GM | MT.PM |
|---|---|---|---|---|---|
| CR | | 0.8466 | 0.8673 | 0.9011 | 0.8762 |
| No_Md | 0 | **6.80E-17** | **2.57E-11** | **3.08E-09** | **6.86E-14** |
| MT.AL | 8.350497 | 0 | 5.71E-01 | 1.13E-01 | 2.37E-01 |
| MT.GP | 6.669175 | 0.5667709 | 0 | 3.52E-01 | 6.07E-01 |
| MT.GM | 5.9274 | 1.5832396 | 0.9300734 | 0 | 6.21E-01 |
| MT.PM | 7.490543 | 1.1830142 | 0.5137082 | 0.4941889 | 0 |

| Batoids | | | | | |
|---|---|---|---|---|---|
| CR | | 0.852 | 0.8729 | 0.9057 | 0.8779 |
| No_Md | 0 | **1.66E-17** | **1.42E-11** | **2.57E-09** | **1.46E-14** |
| MT.AL | 8.515617 | 0 | 5.28E-01 | 9.22E-02 | 2.68E-01 |
| MT.GP | 6.756021 | 0.6310762 | 0 | 3.35E-01 | 7.02E-01 |
| MT.GM | 5.957238 | 1.6838353 | 0.9638557 | 0 | 5.07E-01 |
| MT.PM | 7.691361 | 1.1081814 | 0.3823374 | 0.6640423 | 0 |

### Supplementary Figures

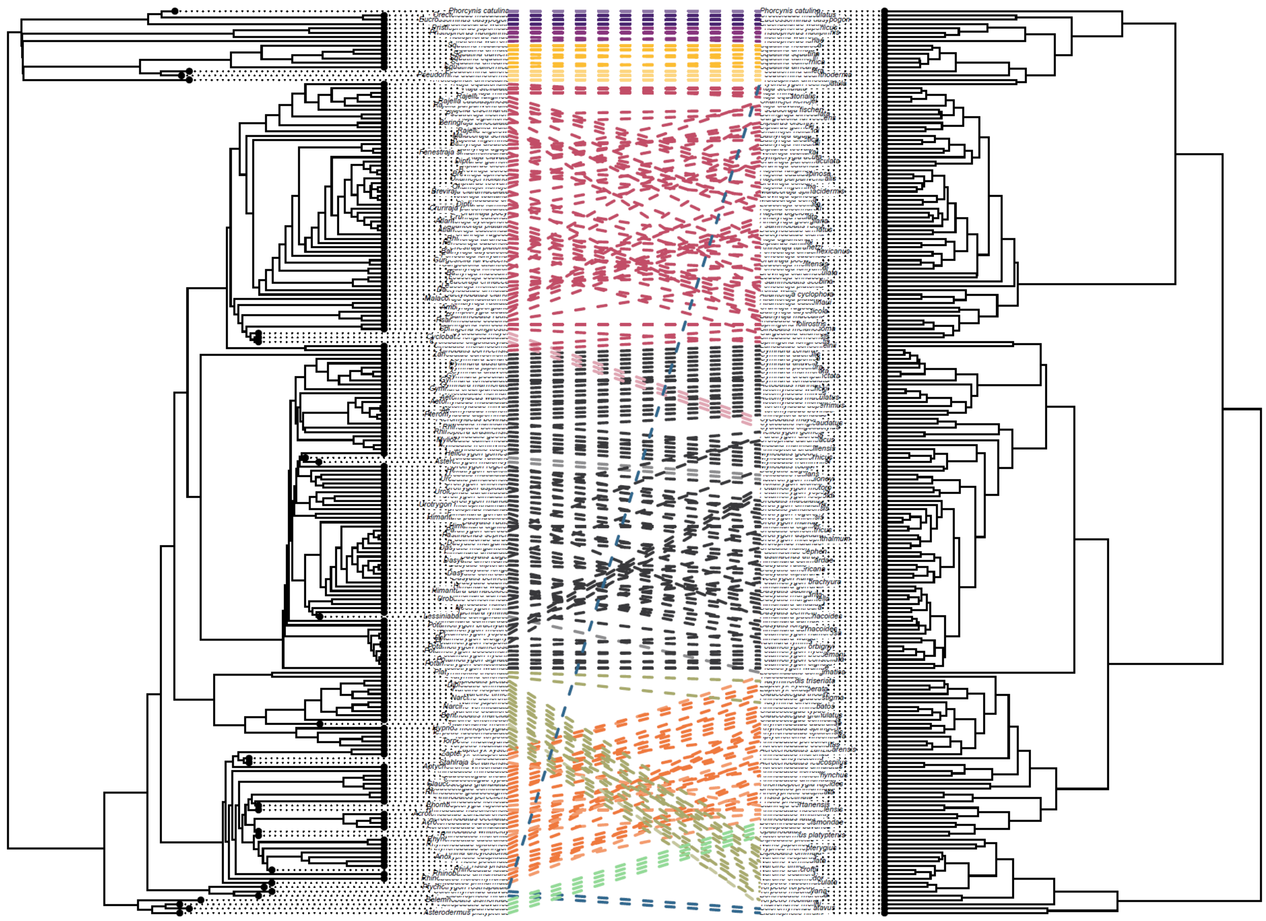

**Supplementary Figure 1.** Phylogenetic tree (left) and distance-based tree (right) using the pairwise procrustes distances matrix followed by a UPGMA clustering. Lines indicate the taxonomic group.

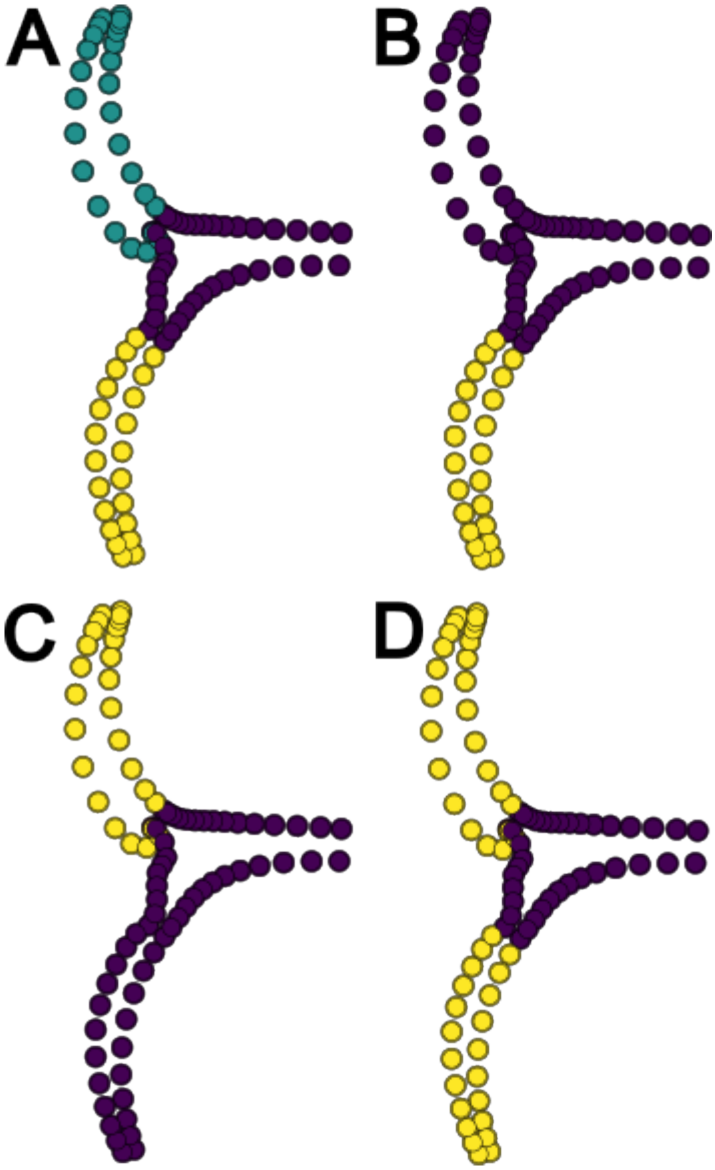

**Supplementary figure 2.** Tested modularity hypotheses. A) Three modules with each element separated. B) Two modules with propterygium and coracoid bar as a single module. C) Two modules as the metapterygium and coracoid bar as a single module. D) Two modules with the propterygium and metapterygium as a single module.

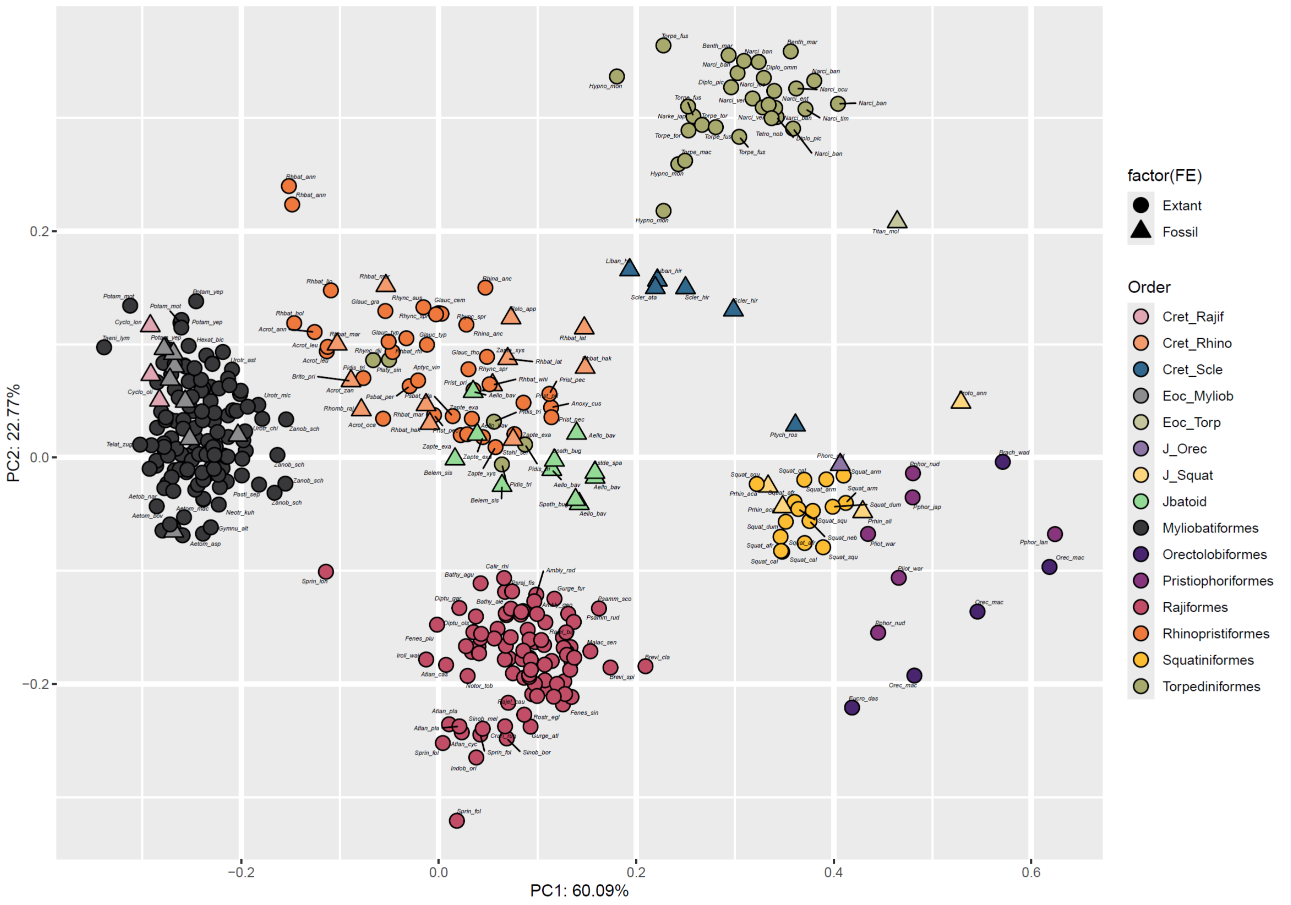

**Supplementary Figure 3.** Principal component analysis with all the individuals used in the study considering the full landmark configuration. Colour code indicate the taxonomic group.

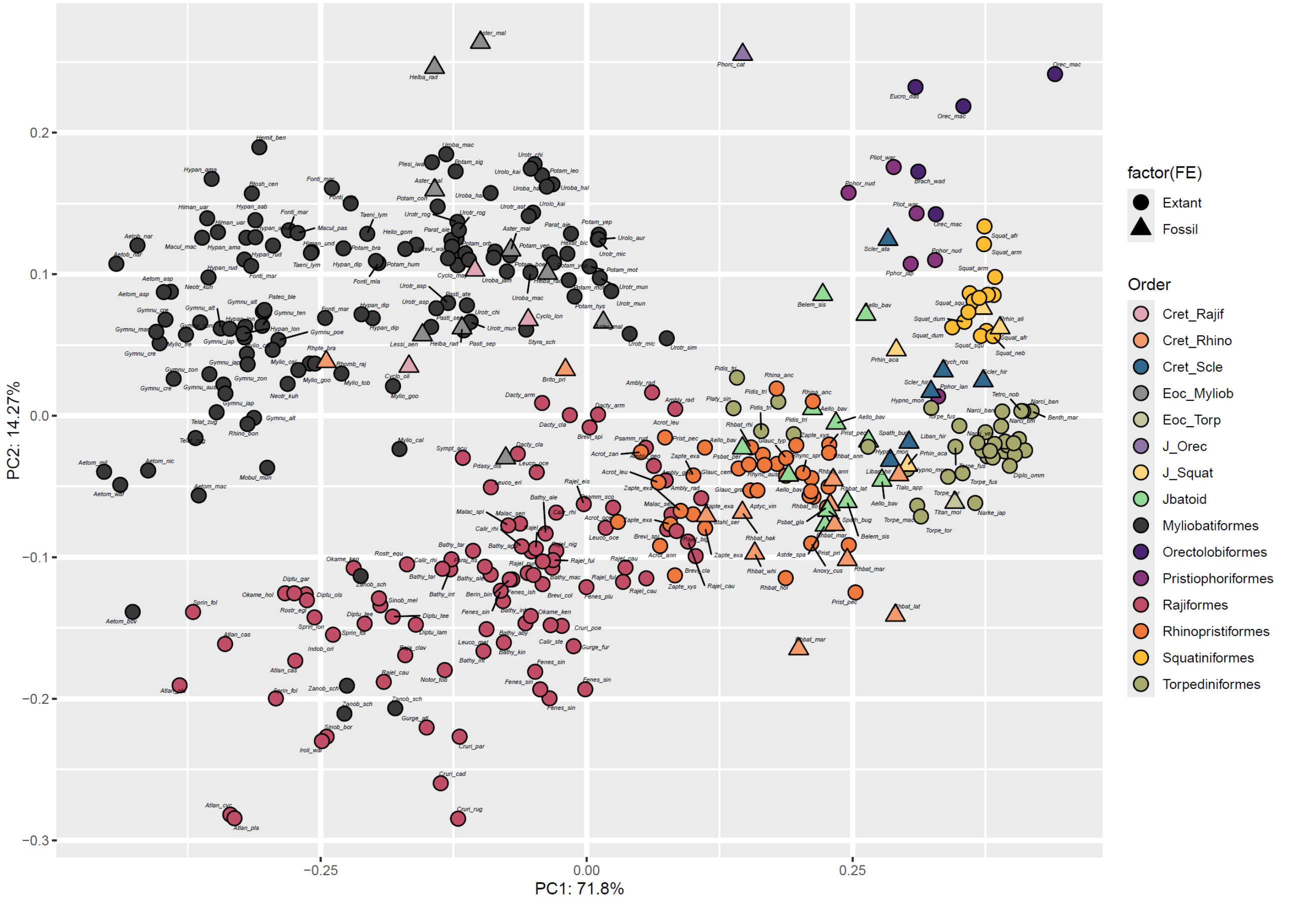

**Supplementary figure 4.** Principal component analysis of all the individuals with only the coracoid bar landmark configuration. Colour code indicate the taxonomic group.

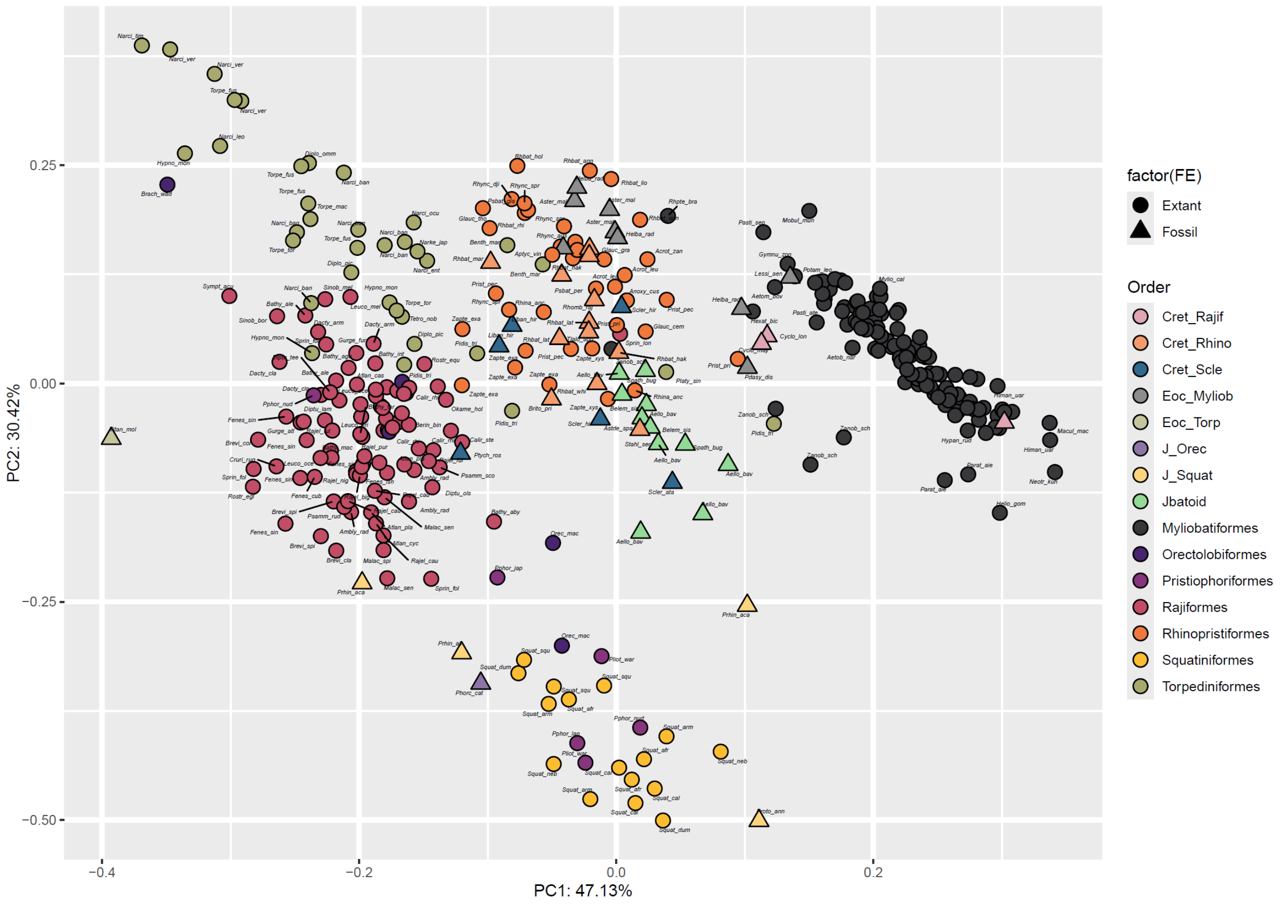

**Supplementary figure 5.** Principal component analysis with all the individuals for the propterygium landmark configuration. Colour code indicates the taxonomic group.

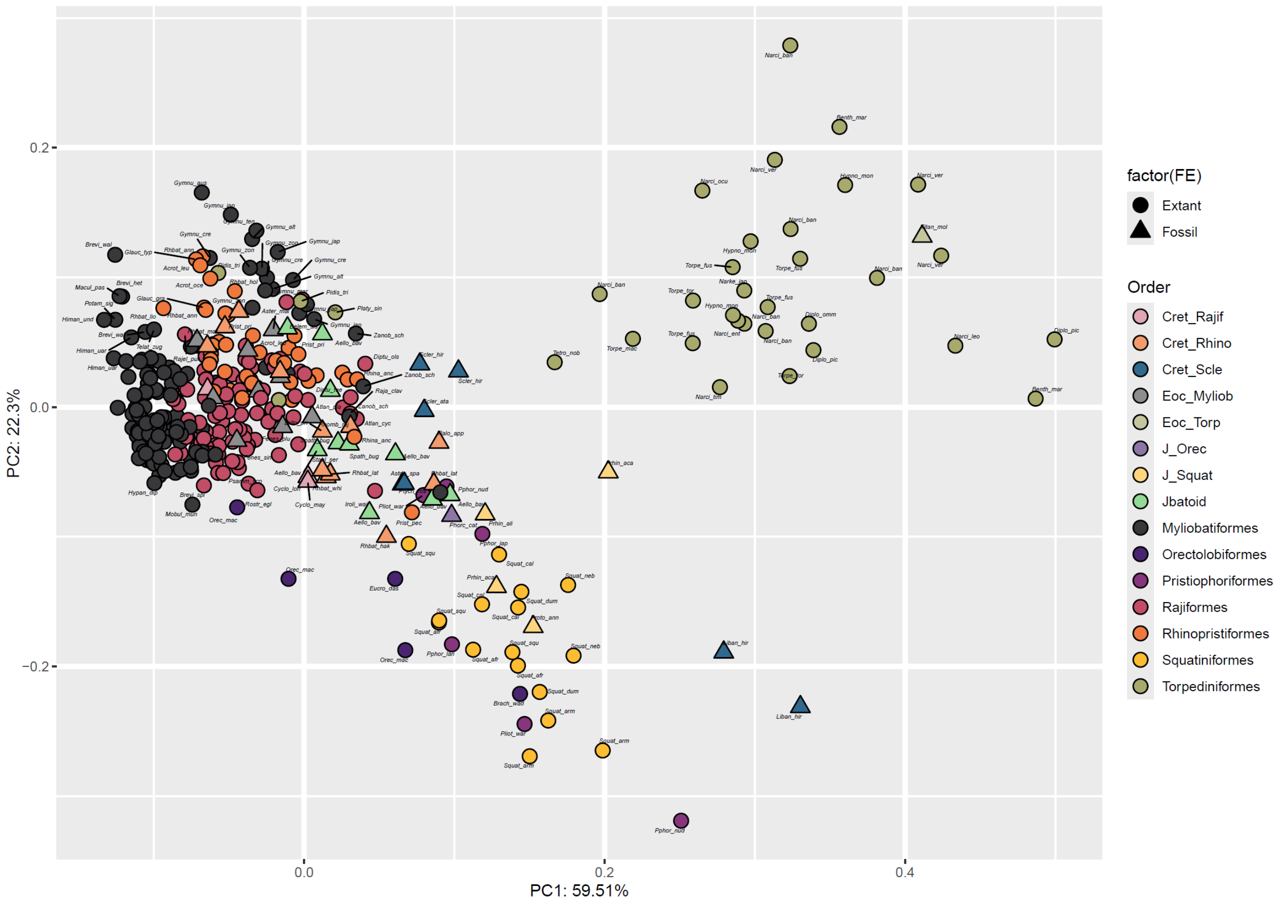

**Supplementary figure 6.** Principal component analysis with all the individuals for the metapterygium landmark configuration. Colour code indicates the taxonomic group.

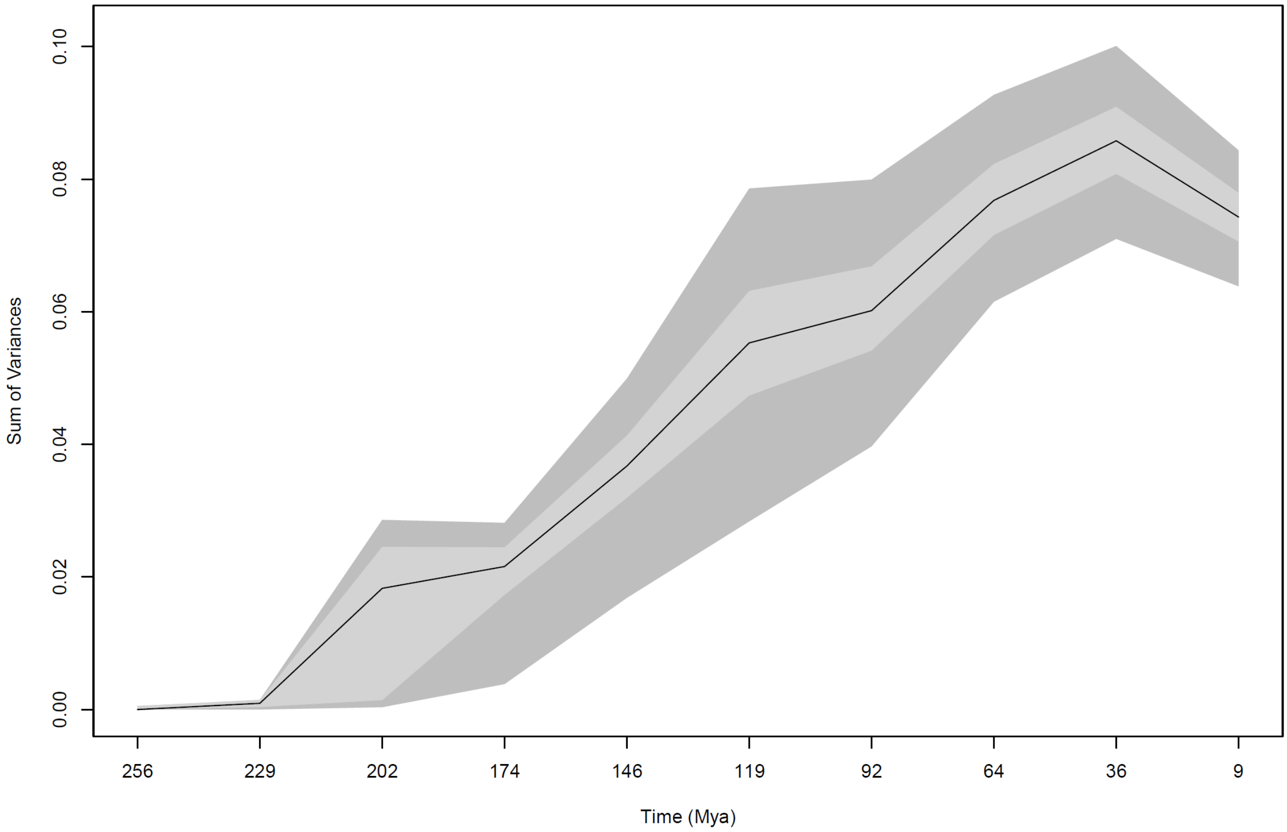

**Supplementary figure 7.** Disparity through time for all the individuals, including sharks and batoids. Disparity estimated as sum of variances.

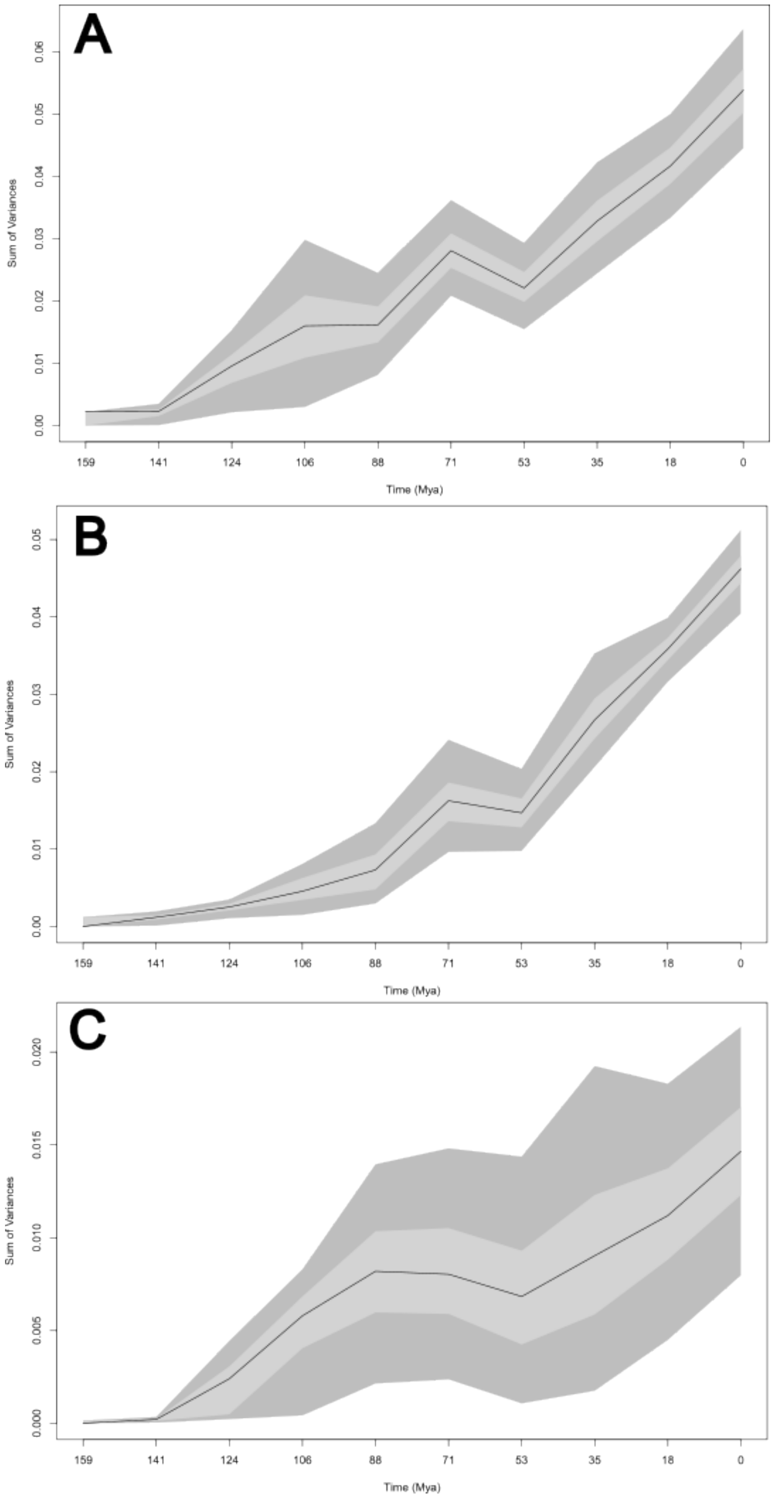

**Supplementary Figure 8.** Disparity through time for only batoid species. A) Coracoid bar landmark configuration. B) Propterygium landmark configuration. C) Metapterygium landmark configuration.

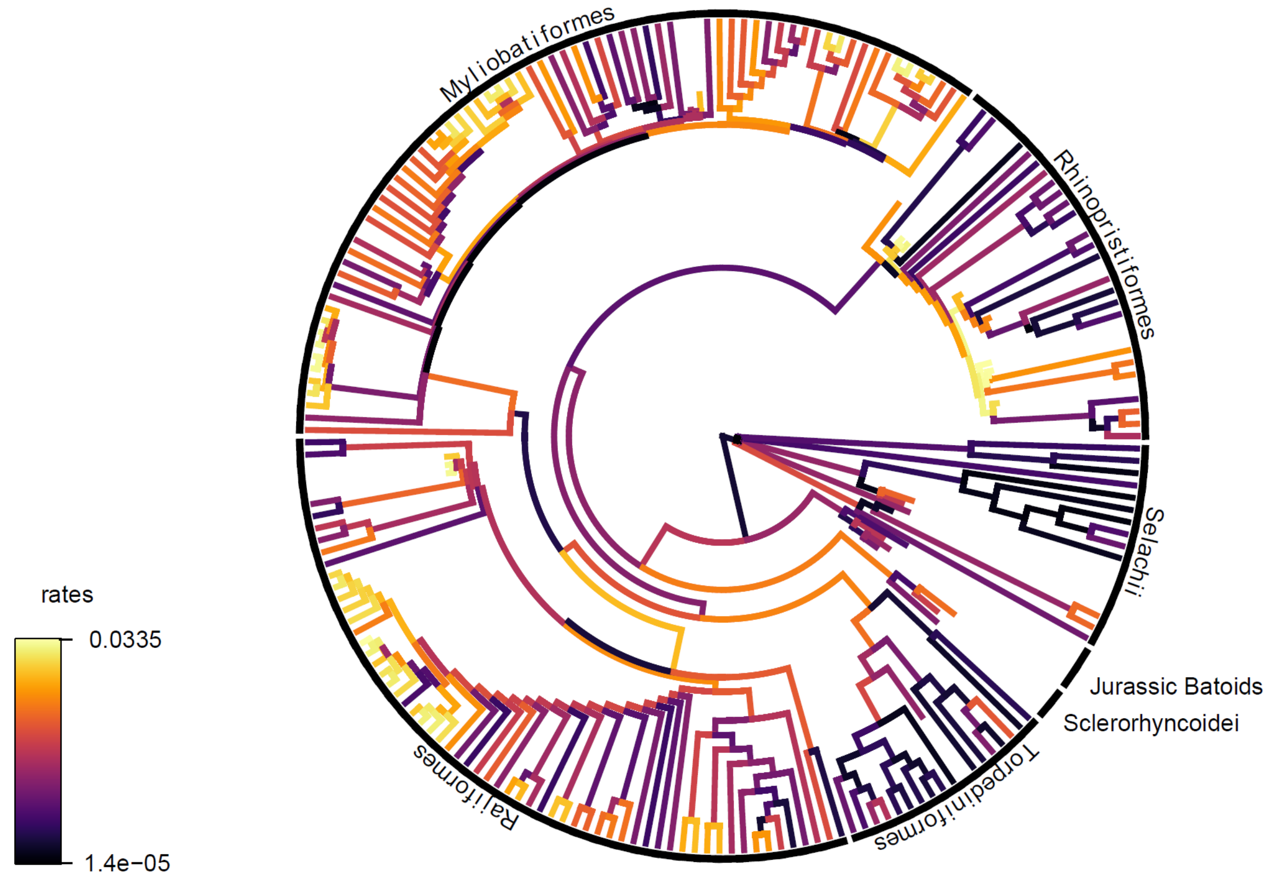

**Supplementary figure 9.** Evolutionary rates per branch for the coracoid bar configuration. Warmer colours indicate higher rates.

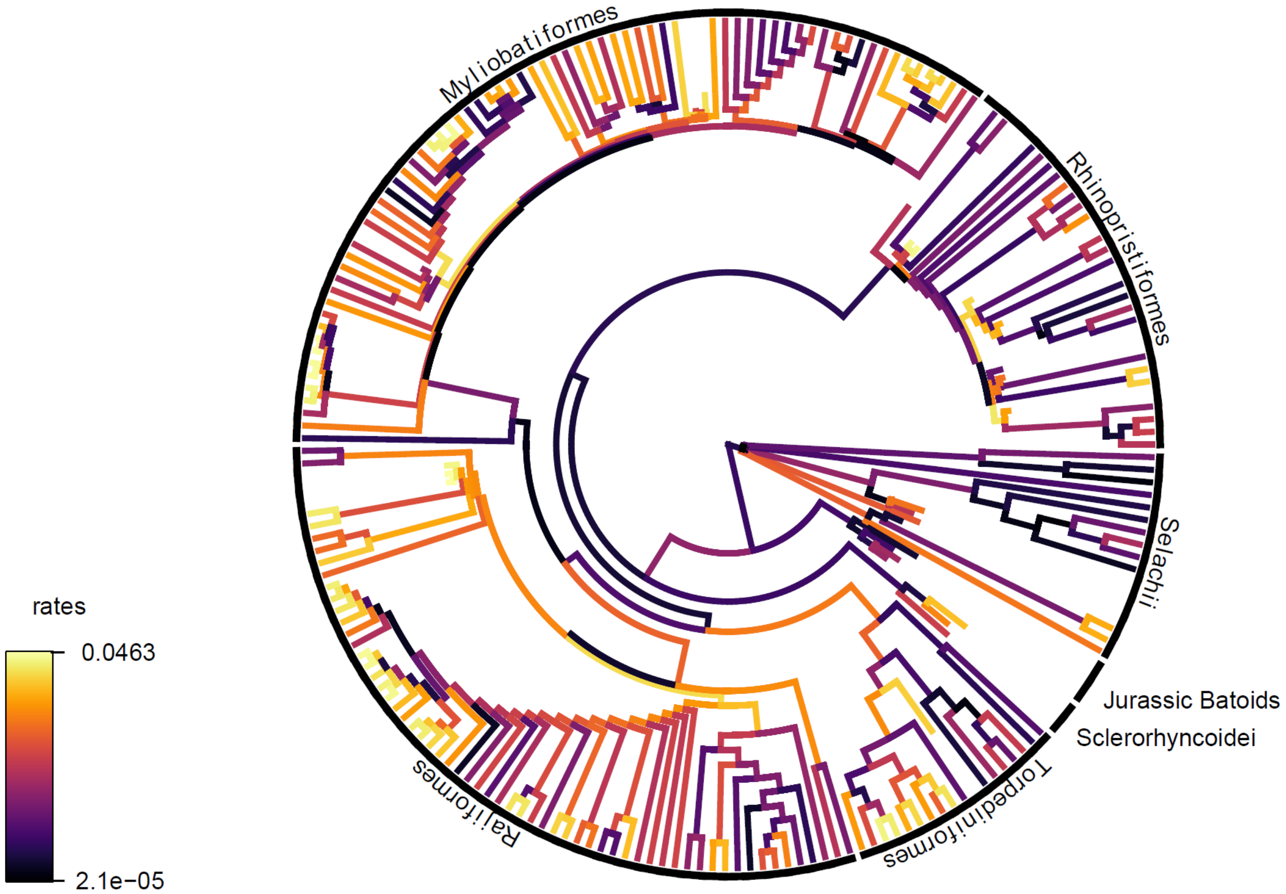

**Supplementary figure 10.** Evolutionary rates per branch for the propterygium configuration. Warmer colours indicate higher rates.

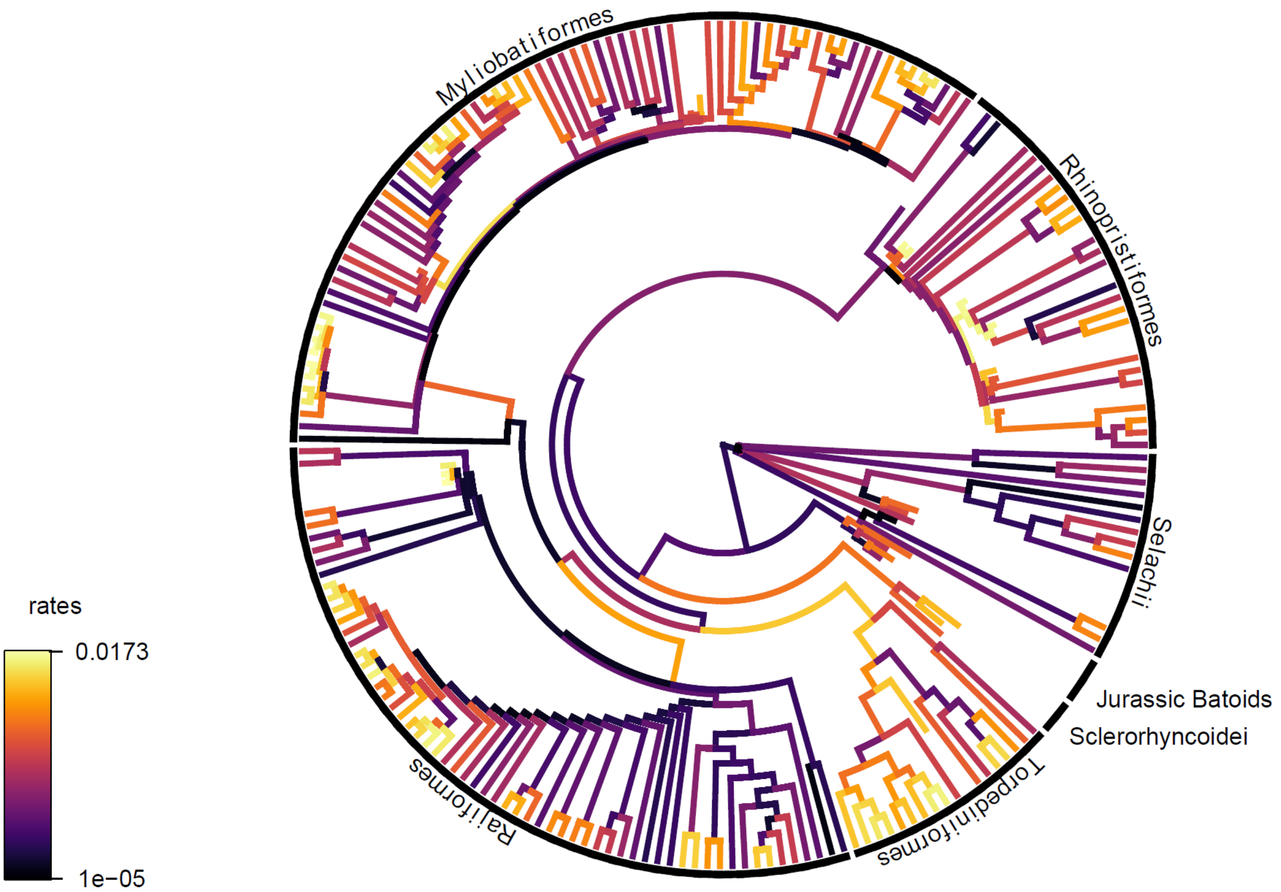

**Supplementary figure 11.** Evolutionary rates per branch for the propterygium configuration. Warmer colours indicate higher rates.
